# The Role Of Circle Of Willis Anatomy Variations In Cardio-embolic Stroke - A Patient-specific Simulation Based Study

**DOI:** 10.1101/190579

**Authors:** Debanjan Mukherjee, Neel D. Jani, Jared Narvid, Shawn C. Shadden

## Abstract

We describe a patient-specific simulation based investigation on the role of Circle of Willis anatomy in cardioembolic stroke. Our simulation framework consists of medical image-driven modeling of patient anatomy including the Circle, 3D blood flow simulation through patient vasculature, embolus transport modeling using a discrete particle dynamics technique, and a sampling based approach to incorporate parametric variations. A total of 24 (four patients and six Circle anatomies including the complete Circle) models were considered, with cardiogenic emboli of varying sizes and compositions released virtually and tracked to compute distribution to the brain. The results establish that Circle anatomical variations significantly influence embolus distribution to the six major cerebral arteries. Embolus distribution to MCA territory is found to be least sensitive to the influence of anatomical variations. For varying Circle topologies, differences in flow through cervical vasculature are observed. This incoming flow is recruited differently across the communicating arteries of the Circle for varying anastomoses. Emboli interact with the routed flow, and can undergo significant traversal across the Circle arterial segments, depending upon their inertia and density ratio with respect to blood. This interaction drives the underlying biomechanics of embolus transport across the Circle, explaining how Circle anatomy influences embolism risk.

## 1 Introduction

The Circle of Willis (hereafter referred to as CoW) is a ring-like network of arteries in the brain, which conjoins the six major cerebral arteries, the internal carotid arteries, and the basilar artery. This network of vessels play a key role in maintaining cerebral blood flow distribution as flow enters from the internal carotids and the vertebro-basilar arteries, enabling collateral flow between hemispheres, and providing a compensatory collateral flow mechanism in case of flow disruption in one of the major supplying arteries [40, 19, 27]. The anatomy of the CoW presents considerable variations among subjects, as documented in several studies [2, 4, 16]. Only about 40% of the population has a well-formed complete CoW anatomy while the remaining manifest some anatomical variations. The dominant anatomical variations are due to under-developed (hypoplastic) or absent (aplastic) artery segments, while fused vessels and additional vessel segments are also sometimes observed [4].

Given its key role in distributing blood and maintaining cerebral perfusion, the connection between CoW anatomy and ischemic stroke is of considerable interest. Prior studies have indicated that the proximal collateral circulation capabilities of the CoW plays an important role in the pathophysiology of cerebral ischemia [19, 29]. The anatomy of the CoW has also been linked to stroke risks in patients with occlusive internal carotid artery disease via medical imaging studies [41, 25, 21, 19, 20, 49]. For example, trans-cranial Doppler imaging in conjunction with carotid compression has been employed to characterize CoW collateral capability in [21, 19, 20], while MR imaging on patient population with reported stroke incidences have been used to explore the link between ischemia and CoW anatomy in [41, 16, 49]. Majority of these investigations are clinical, based on imaging studies on patient cohorts, and the insights presented here are primarily based on compensatory hemodynamic flow through the communicating arteries.

It is well-known that a significant proportion of ischemic strokes and transient ischemic attacks (TIA) are ultimately due to an embolus [10, 6], and CT or MR imaging of the occlusion location often fails to provide information on the source of the embolus causing the ischemia (leading to difficulty in diagnosis in the presence of multiple potential sources of embolisms) [11, 15]. This motivates investigating how CoW anatomy impacts cerebral embolic events. However, this has remained a major challenge since it is difficult to control for the anatomy of the CoW and model embolus dynamics in patient-cohort studies or animal models. Recent advances in medical image-driven computational modeling have opened up the possibility of addressing this challenge via systematic *in silico* studies on image-based vascular models[43, 46]. Specifically, flow in the CoW, and the influence of anatomical variations on hemodynamics, has been studied using one-dimensional reduced-order network models [17, 18, 1, 9], three-dimensional computational fluid dynamics simulations on representative CoW geometry [3], and hybrid techniques [14].

In a series of prior works [7, 36, 35, 37], we have established a computational pipeline for studying embolus transport within 3D patient-specific hemodynamics. Our computational model comprises: (a) medical image-based modeling of vascular anatomy; (b) fully resolved 3D time-dependent flow simulations; (c) discrete particle method for embolus transport; and (d) a Monte-Carlo approach for characterizing embolus distribution statistics. We have addressed mathematical modeling issues pertaining to embolus transport [36, 37], elucidated how embolus size and other factors influence embolus distribution to the brain [7, 35], characterized differences between the distribution of cardiogenic and aortogenic embolus distribution to the brain [35], and demonstrated that understanding the chaotic advection of emboli through large arteries is necessary for discerning the source-destination relationship for cerebral emboli [37]. In this work, we investigate the link between CoW anatomical variations and transport of emboli, with two specific objectives. The first objective is to test the hypothesis that the anatomy of CoW does influence embolus distribution in the brain. The second, is to describe a mechanistic reasoning for this influence based on hemodynamics.

## 2 Materials and Methods

### 2.1 Selection of anatomical variations for Circle of Willis

A broad literature search was performed for research articles comprising population data on occurrence frequencies of various CoW anatomical variations. A set of 25 articles *(reference list included in supplementary material)* were selected from the search results. Occcurrence frequencies of the various commonly observed CoW anatomical variants were extracted from these articles. Using ranking statistics on this data, a list of anatomical variations arranged in order of frequency was obtained. The 5 most frequent variations were chosen for this study. These involved an absent/hypoplastic anterior communicating artery (AcoA), left and right posterior communicating arteries (L.Comm, R.Comm), and left and right P1 (LP1, RP1) connectors (fetal type posterior cerebral artery anatomy). The respective complete CoW anatomy was also considered - resulting in a total of 6 anatomical variations of the CoW being studied per patient. Schematic representation of these anatomical variants have been presented on the top-row in Figure 1, along with labels for the various vessels, which will be consistently employed throughout this paper. Namely, middle, anterior, and posterior cerebral arteries will be referred to as MCA, ACA, and PCA respectively. Internal carotid and basilar arteries will be referred to as ICA and BA respectively. The suffix R/L denotes right/left.

**Figure 1:**
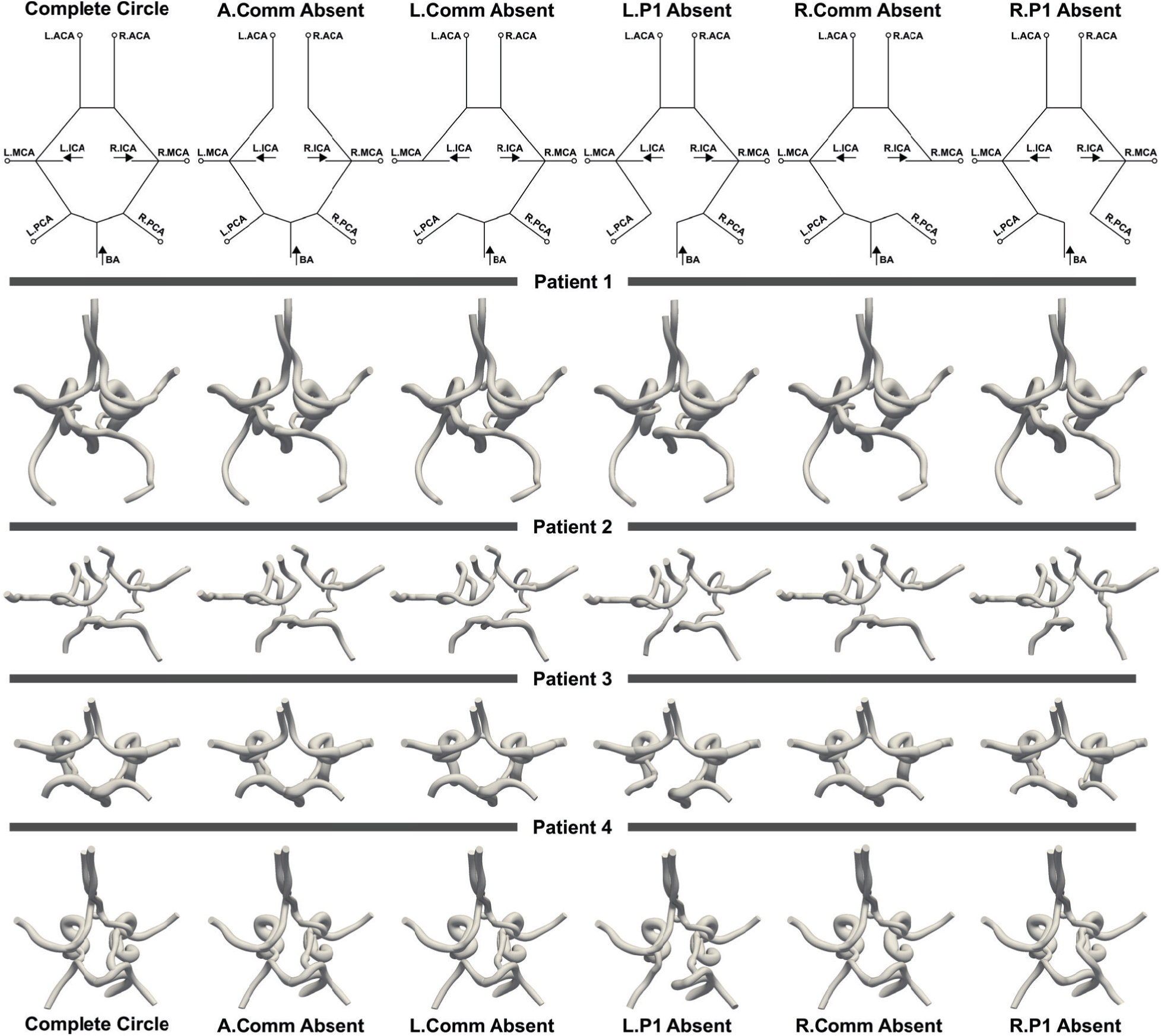
An illustration of all the 24 different Circle of Willis anatomical models considered for this study. Note that the zoom and scale of each has been adjusted so that all of them fit the same width on this image-panel. Top row depicts schematic versions of the respective anatomical variant, with artery labels followed in the study. L/R denote left and right; ACA, MCA, PCA denote anterior, middle, and posterior cerebral arteries respectively; ICA is the internal carotid artery, and BA denotes the basilar artery.

### 2.2 Image-based modeling of anatomy

A set of four representative patient geometries, used in our prior works [35, 37], was selected for this study, from amongst a larger set of computed tomography (CT) images acquired as a part of the Institutional Review Board (IRB) approved Screening Technology and Outcome Project in Stroke (STOP-Stroke) database study [42]. The patient anatomies were selected such that they possessed a complete CoW, and well differentiated branching vessels at the aortic arch (that is, no fused vessels or bovine arch anatomy). Artery lumen segmentations were generated from patient CT images using the custom 2D lofted image-segmentation technique incorporated in the open-source cardiovascular modeling package SimVascular [48]. During segmentation, it was ensured that cerebral vasculature up to the M1, A1, and P1 segments of the MCA, ACA, and PCA respectively, were included for each patient. The image segmentations were used to generate 3D computer solid models of the complete arterial pathway from the aorta to the CoW. In order to incorporate anatomical variations of the CoW within this image-based modeling procedure, respective communicating artery segmentations were iteratively detracted from the original segmented data, and the modified segmentations were re-lofted to generate 3D models of the corresponding incomplete CoW anastomoses. By virtually introducing CoW variations in a given patient model (as opposed to a patient with that anatomical variant inherent) we could control against other inter-patient variabilities, and isolate the effect of CoW anatomy. Spanning across four patients, each with a total of six CoW topologies described in Section 2.1, a set of 24 anatomic models were thus created, which have been illustrated in Figure 1. Each model was discretized into a computational mesh for performing finite-element based hemodynamics simulations using SimVascular [48]. The average number of linear tetrahedral elements for each computational model ranged between 6.0-8.5 million, with average element size ranging between 0.36 mm and 0.52 mm, a choice that was guided by detailed mesh refinement and flow analysis studies as described in [28]. Further details on the modeling and meshing operations, and mesh refinement, have been discussed in prior studies [36, 37].

### 2.3 Hemodynamics simulation

For hemodynamics simulations, blood was assumed to be a Newtonian fluid, with bulk density of 1.06 g/cc, and viscosity of 4.0 cP. Three-dimensional blood flow through each of the 24 anatomical models was computed using the classical Navier-Stokes equations and continuity equation, describing momentum and mass balance, respectively, for blood flow. A Petrov-Galerkin stabilized linear finite-element method was employed to solve these equations for calculating flow velocity and pressure fields. The summary of equations, including the variational form for finite-element method, has been presented as follows:

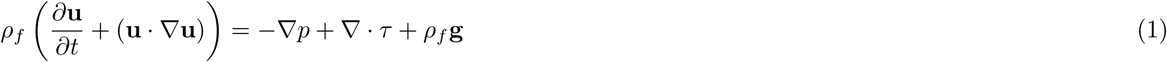

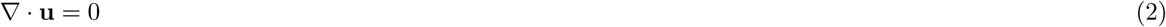

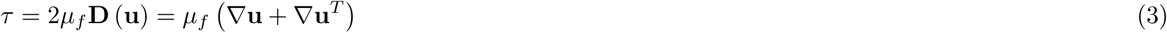

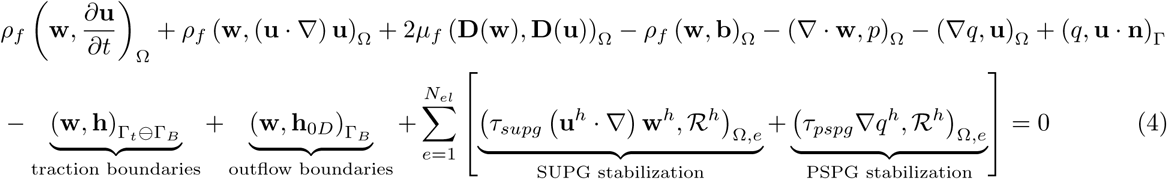

where Eq. (1) is the Navier-Stokes equation, Eq. (2) is the continuity equation, Eq. (3) describes the constitutive stress-strain law for Newtonian fluids, and Eq. (4) describes the complete, generalized variational form of Eqs. (1)-(3) for the stabilized finite element method employed here. In Eqs. (1) - (4), **u**, *p* denote the blood flow velocity and pressure, **w**, *q* denote the finite element test functions, *µ*_*f*_ is the blood viscosity, *ρ*_*f*_ is the blood density, **g** denotes gravity, Ω indicates the overall computational domain encompassing the arterial network, Γ and its sub-scripts denote the various boundary faces of the computational domain, 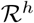 denotes the residual of the momentum equation, and finally *τ*_*supg/pspg*_ are stabilization factors for the SUPG stabilization (for convective flow) and PSPG stabilization (for pressure stability using equal order finite elements) terms respectively. A common pulsatile flow profile, originally derived from magnetic resonance measurements [38], was imposed at the inlet of the model at the aortic root to drive the flow. The average volumetric inflow was kept consistent across all the models at 79 ml/s (4.8 l/min). At the outlet boundaries, the influence of the downstream vascular beds were incorporated by assigning resistance based boundary conditions (that is, **h**_0*D*_ in Eq. (4)), which couples the outlet pressure with computed outlet flow rate. Resistance values were assigned by estimating a total arterial resistance and distributing this resistance across the outflow boundaries based on target flow division across the outlets [35]. Target ranges were set in three steps. First, 65% of total flow was assumed to exit the descending aorta [7]. Second, target ranges for the six cerebral arteries were assumed based on measured MR data across 30 patients as reported in [31]. Finally, remaining flow was assigned to subclavian and external carotid outlets based on their respective cross-sectional areas [50]. Once the preliminary estimates of resistance values were assigned, steady flow simulations were run for each model to iteratively tune these values to ensure that average outflow for each outlet was within the assumed target ranges [35, 37]. Specifically in the present study, the target flow ranges for this procedure, for the six cerebral arteries, were held the same for all the 24 patient-CoW combinations. Additional numerical details for these steps have been extensively discussed in [7, 35, 37].

### 2.4 Embolus transport

Emboli were assumed to be spherical particles, and their trajectory across the arterial pathway were computed using a one-way coupled scheme, which assumes that a particle is influenced by the flow, but not vice-versa, in the large arteries modeled here. The fluid-particle interaction was based on a custom modified form of the classical Maxey-Riley equation [33]. Our modifications entailed incorporating near-wall shear gradient driven lift forces, accounting for particle collisions with artery walls, and elastohydrodynamic lubrication effects near-wall. The final form of the resultant modified Maxey-Riley equation is summarized as follows:

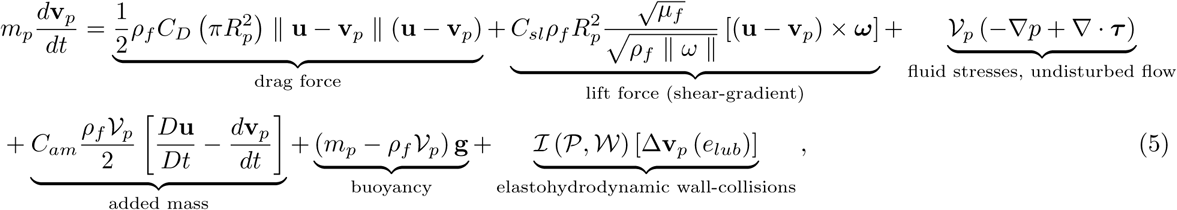

where *m_p_* denotes the mass of the particle (embolus), 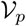 is the particle volume, ***v**_p_* is the particle translational velocity, and *R_p_* is the particle radius. The terms *C*_*D*_ and *C*_*sl*_ denote the drag and shear-gradient lift force coefficients, computed using expressions described in [37]. Similarly, *C*_*am*_ denotes the added-mass coefficient, which is unity for spherical immersed bodies. The term *ω* = *∇ ×* **u** denote the flow vorticity at the location of the particle. The function 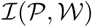 is an indicator function, assuming a value of 1 if the embolus particle 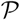 contacts the vessel wall 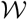, and 0 otherwise. The term [∆**v**_*p*_ (*e*_*lub*_)] represents the contribution to momentum change induced by elastohydrodynamic lubrication and particle-wall collisions, modeled using a lubrication dependent restitution coefficient *e*_*lub*_. The respective mathematical expressions and algorithmic issues for each of these terms have been described in more detail in [37, 35, 36]. Each individual particle trajectory was integrated in time using an explicit Euler time integration scheme with a time-step of 0.1 ms.

### 2.5 Sampling based statistical framework

The combined fluid-particle scheme was embedded within a Monte Carlo sampling based framework, developed in [35]. A multi-parameter design of experiments was devised, by considering two different embolus material compositions (labelled as 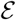), and (for each material) three different embolus sizes (labeled as 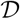). Ensembles of spherical emboli were generated at locations along the aorta inlet face, representing cardiogenic emboli entering the aortic arch. Each ensemble comprised approximately 4,020 particles. Eight characteristic release instances were chosen across the cardiac cycle (labeled as 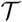), where embolus samples were released into the flow, and their trajectories were integrated. The overall list of parametric combinations have been compiled in Table 1. For each parameter combination, particle distribution to each vessel outlet was computed over 10 cardiac cycles. Combined simulation runtime for the flow and embolus simulations for each model was around 24-36 hours on an average of 80 cores on the Berkeley Research Cluster supercomuter. Following the notation described in [35], the number fraction for the *k*’th vessel is defined as *φ_k_*(Λ) = *N_k_/N*_*s*_, where *N*_*k*_ is the number of embolic particles reaching the vessel, *N*_*s*_ is the total number of particles in the ensemble, and Λ denotes the input vector comprising a combination of the various parameters defined here i.e. 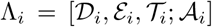, where 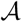 is a place-holder for the anatomy of the CoW (Table 1). We further processed these number fractions *φ*_*k*_ to account for combined effect of underlying variables as well as to denote collections of vessels. For example, *φ*_*cow*_ will denote the total number fraction reaching the CoW, 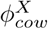 will denote summation of number fractions across all variable combinations for the *X*’th variable (i.e. 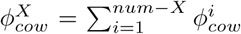). For the present study, a variable of specific interest is the ratio 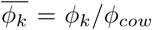 for each of the six cerebral arteries, which denotes the proportion of cerebral emboli reaching the *k*’th cerebral artery, and sums to a total of 1.0 across all the six cerebral arteries.

**Table 1:**
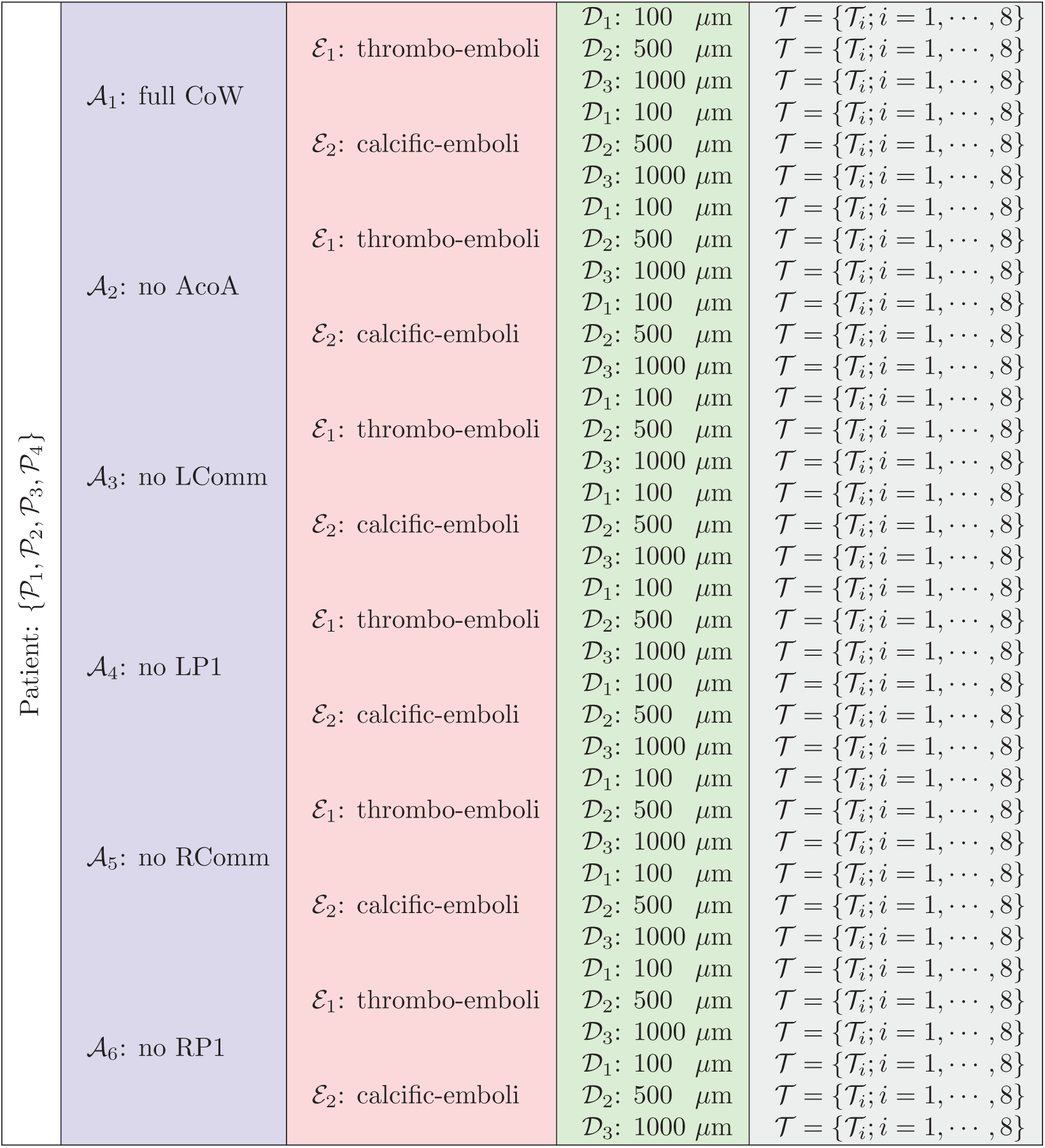
Table outlining the various parameter combinations incorporated in the simulation study presented here. The full set of numerical experiments presented here spans a total of 1152 parameter combinations, with around 4,020 virtual embolus particles tracked per combination.

The computed number fractions were employed to establish whether CoW anatomy has a significant, and quantifiable, influence on embolus distribution in the brain. This was achieved by computing variabilities induced in embolus distribution to cerebral arteries with respect to CoW anatomy, over and above any variabilities expected from flow distribution alone. The distribution fraction 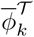 was computed for the six cerebral arteries. For each embolus composition and size, this fraction was compared across all CoW anatomical variants. Thereafter, the sample mean and standard deviation of the respective distribution fractions 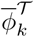 were estimated. A relative measure of variability in embolus distribution fractions for each cerebral artery was obtained by computing the coefficient of variation (*cov_e_*) - defined as the ratio between the standard deviation and the mean of the distribution fraction data (i.e. 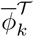 for various parameter and anatomy combinations 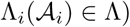. Additionally, for the varying CoW anatomies, the corresponding mean, standard deviation, and coefficient of variation (*cov*_*f*_) for the flow distribution ratios (i.e. similar to *φ*_*k*_ above, 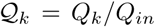 *etc.*; *Q* denoting volumetric flow, *Q*_*in*_ total cardiac output) to each cerebral artery were computed. Dividing *cov*_*e*_ by the corresponding *cov*_*f*_ values ensured that the variability in embolus distribution is normalized against any existing variability induced by the flow distribution.

### 2.6 Flow and embolus routing through Circle of Willis

Blood flow entering the CoW via the ICAs and the BA is redistributed across the CoW artery network to meet the flow demands for each of the six cerebral artery territories. This flow routing across the communicating segments of the CoW is crucial for determining the proximal collateral capability of cerebral vasculature. While hemodynamic forces drive embolus dynamics across the arterial network, embolus trajectories may not exactly follow the flow routing across the CoW communicating arteries owing to their finite inertia (leading to a finite response time for emboli to influenced by flow-induced forces) and their collisions with vessel walls. To investigate this underlying mechanistic reasoning, and characterize flow routing and embolus routing, a detailed post-processing protocol was devised for the CoW anatomical variants considered. The steps of this procedure have been schematically outlined in Figure 2. First, the seven communicating arteries from the complete CoW anatomy were computationally extracted (Figure 2.a.). Thereafter, the centerlines of the extracted segments were computed, and a set of cross-sectional slices were created normal to the centerline along the length of each of the vessel segments (Figure 2.b). Following this, blood flow velocity fields were interpolated at the locations of these section-planes, and the flow velocity as well as the volumetric flow rate carried by the communicating vessels were computed (Figure 2.c.). Finally, a numeric index for the proportion of cerebral emboli carried by the vessel segments was computed. This was achieved by choosing the midpoint of the centerline of each communicating vessel as a reference point, and identifying points along each embolic particle pathline that lie closest to this reference. Each embolus trajectory was assigned a scalar value equal to the distance between these two points, and the trajectories were clustered based on this distance. For each vessel segment, the smallest coherent cluster size was used as a metric for the total number of emboli carried by the vessel in question (the trajectories marked in blue in Figure 2.d.). This quantity has hereafter been referred to as routing count, denoted by *R*_*k*_ for various communicating artery segments indexed by *k*. We remark that a simple counting of particles across the communicating segments would not be sufficient, since: (a) each particle can potentially travel across several vessel segments as they traverse the circle and exit into one of the six cerebral vascular beds, and (b) particles can often reverse their trajectories under retrograde/swirling flow.

**Figure 2:**
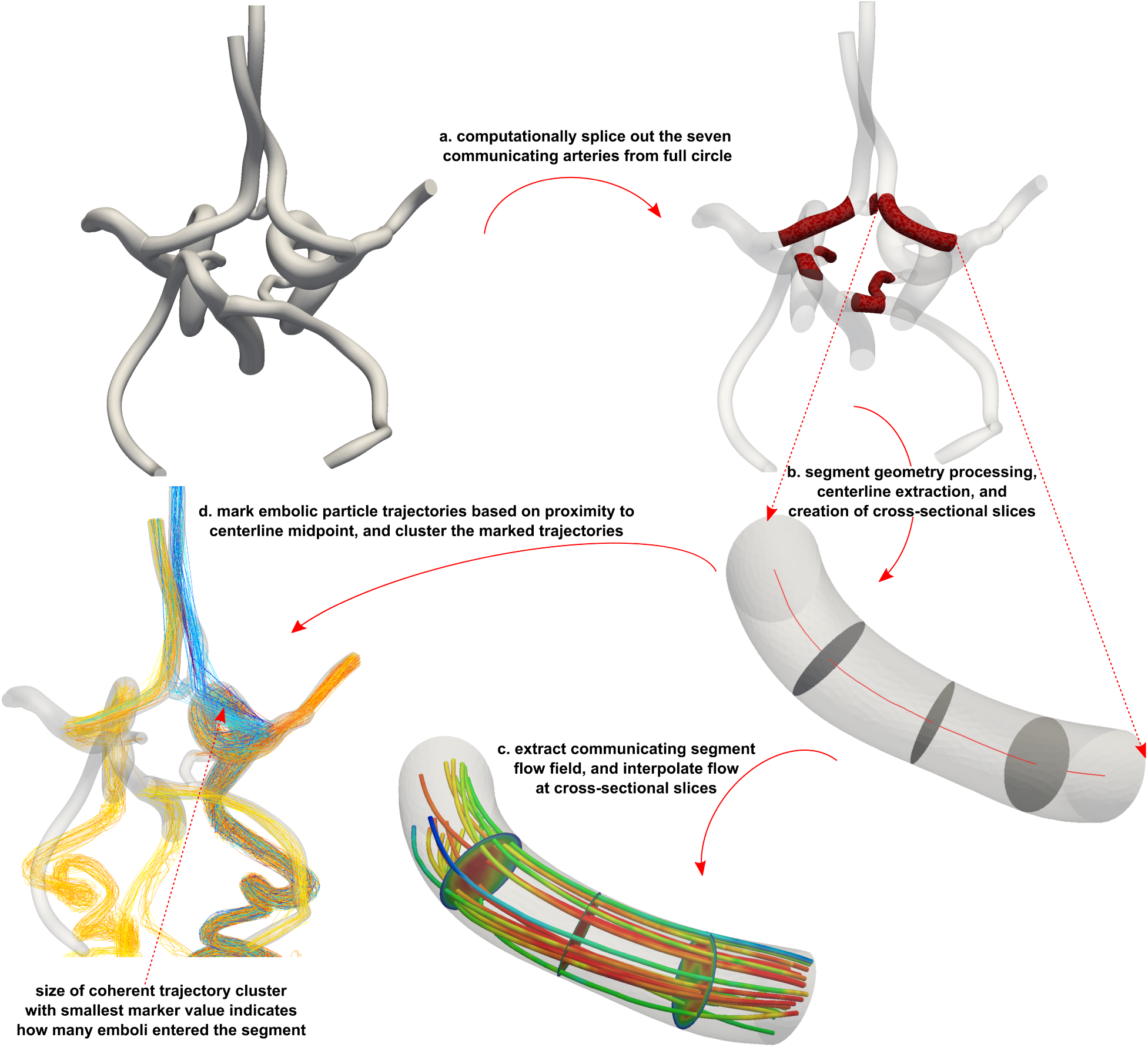
A schematic illustration of the procedure to analyze the routing of flow, and embolus traversal, across the various artery segments of the Circle of Willis.

## 3 Results

### 3.1 Circle of Willis anatomy influences embolus distribution

The ratio between the computed *cov*_*e*_ and *cov*_*f*_, as described in Section 2.5, were compiled for thromboemboli and calcified-emboli of varying sizes, and the sample statistics of these *cov*_*e*_: *cov*_*f*_ ratios, for each cerebral artery, has been illustrated in Figure 3. Table 2 shows the corresponding ratios for each embolus type aggregated over all 24 patient-CoW anatomy combinations. Values equal to unity denote that variabilities in embolus and flow distribution are equivalent, while values greater than unity suggest that embolus distribution manifests greater variability as compared to flow. A value of 1.0 is marked using a green solid line on Figure 3, and a significant number of samples lie above this line, indicating that the respective evaluated ratios deviate from unity, as also observed from Table 2. We employed a non-parametric Wilcoxon test to quantifying how significantly these ratios differ from unity. Corresponding p-values were *<* 0.001 for left and right ACA, left and right PCA, and left MCA, and 0.02 for the right MCA. The particular case of the MCA is further discussed in the next section. These observations indicate that scatter in embolus distribution is substantially different from what we would expect had flow distribution variabilities alone influenced it. This is further illustrated by visualizing the flow and embolus distribution ratios and their variabilities in Figure 4 for thrombo-emboli of varying sizes across all 24 patient-CoW anatomy combinations. Following the description of the parameter combinations in Section 2.5, we note that for each computed *cov*_*e*_ value, embolus properties were held fixed, and release times were summed over, leaving, for each patient, the CoW anatomy as the lone variable. Hence, the observed variabilities in embolus distribution here for each embolus type and size, can be attributed to the influence of anatomical variations *(see also supplementary figure S1)*.

**Table 2:**
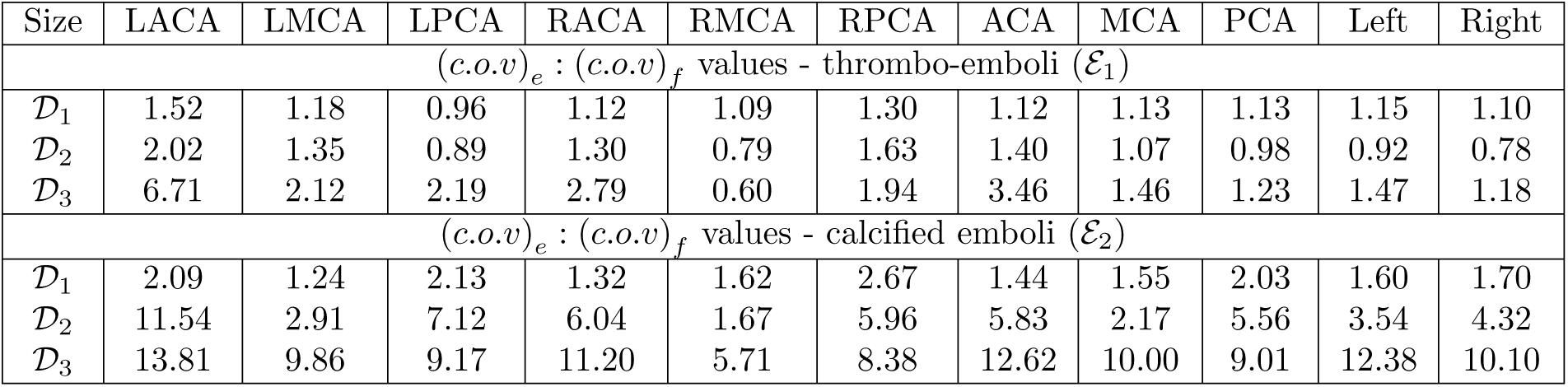
Ratios between embolus and flow coefficients of variation for each cerebral artery, and their combined territories, aggregated across all 24 patient-anatomy combinations. Refer to supplementary tables S1 & S2 for sample data described for each patient.

**Figure 3:**
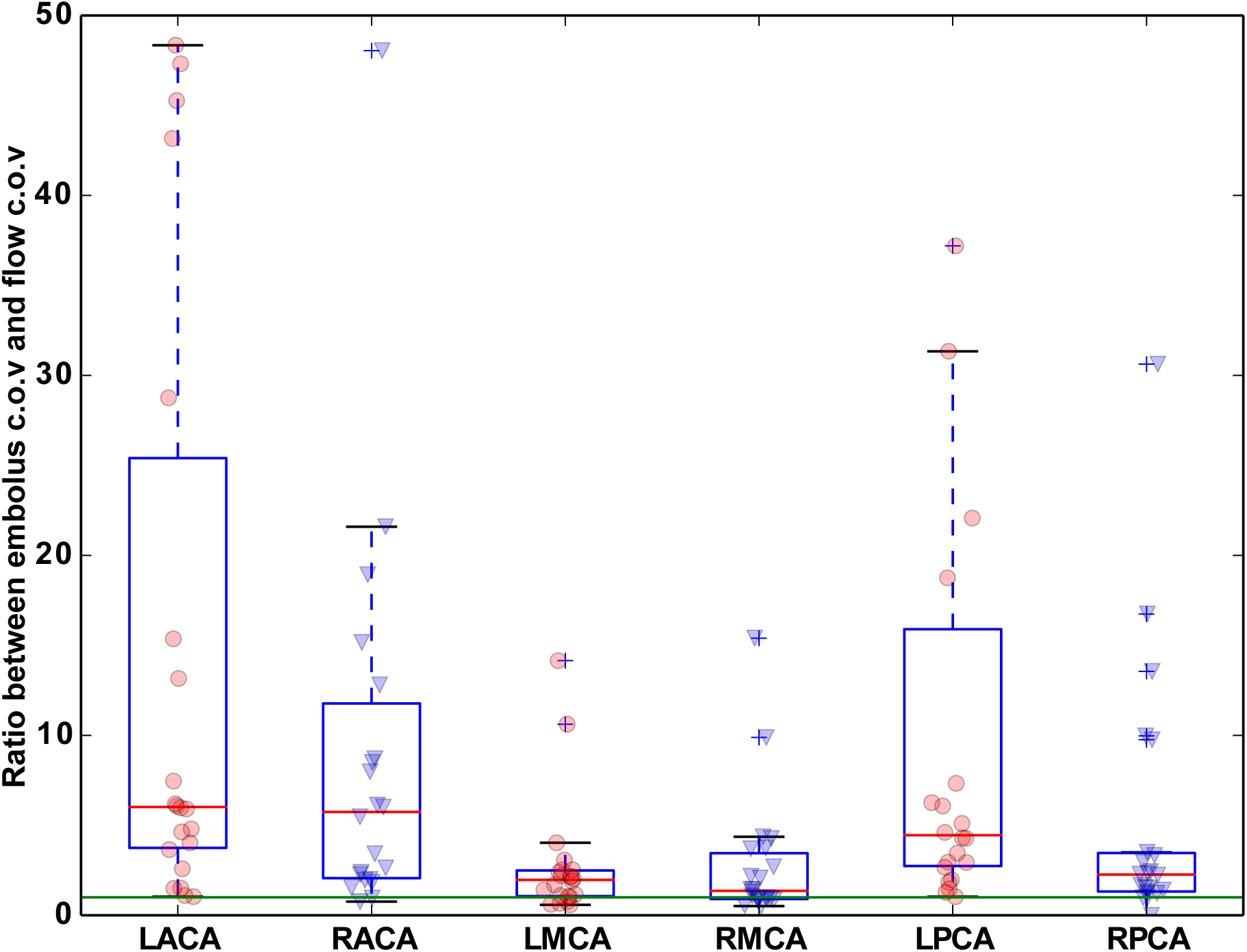
Sample statistics for the ratio between the coefficient of variation for embolus distribution (both thrombo-emboli and calcific emboli) (c.o.v)_e_ w.r.t that for flow distribution (c.o.v)_f_ for each cerebral artery. A ratio of unity (the green solid line) indicates variability induced in embolus and flow distribution are equal, and values greater than unity denote that variability induced in embolus distribution is greater than that expected from flow alone. Supporting sample data for coefficients of variation included in supplementary table S1 & S2

**Figure 4:**
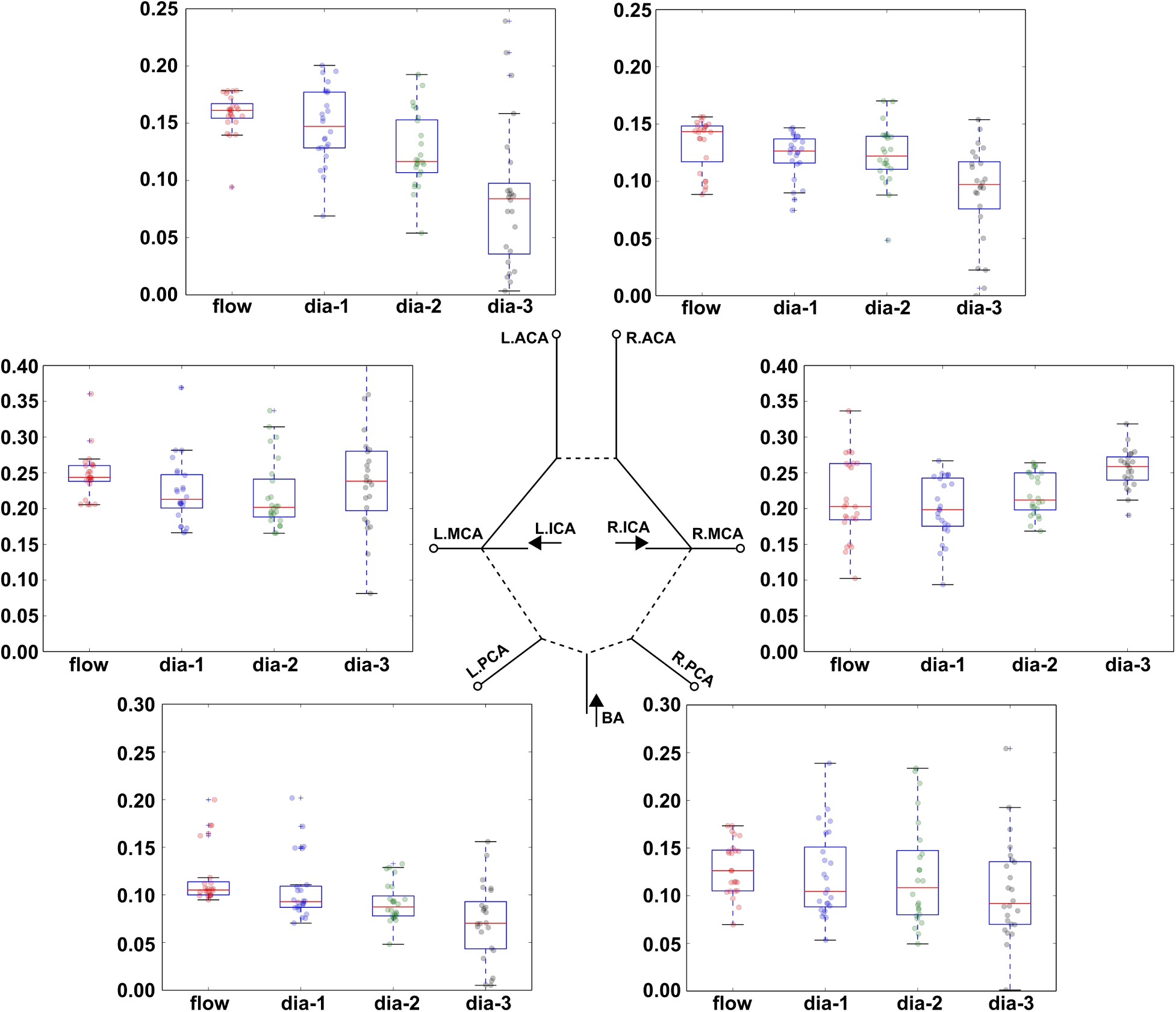
A combined illustration of the distribution fractions of thrombo-emboli to each cerebral artery (plotted along the y-axis of each cerebral artery boxplot), as a function of embolus size, computed across all patient-CoW anatomical models. The corresponding variabilities in flow distribution have also been included. Supporting sample data for coefficient of variation included in supplementary table S1

### 3.2 Embolus distribution to MCA territory is least sensitive to Circle anatomy

The extent of variability in embolus distribution induced due to CoW anatomy variations are different across the six major cerebral arteries. Specifically, from Figure 3 we see that the *cov*_*e*_ values, and the corresponding *cov*_*e*_: *cov*_*f*_ ratios, for both thrombo-emboli and calcified emboli are the lowest for right and left MCA, across all embolus sizes and CoW anatomical variants. Correspondingly, embolus distribution ratios for all sizes considered are greatest for the right and left MCA, as seen in Figure 4 for thrombo-emboli. This indicates that cardiogenic embolus distribution to the MCA territory is least sensitive to potential variabilities induced due to CoW anatomical variations. Figure 3 also indicates that, for the various cases considered here, the variability in cardiogenic embolus distribution to the right cerebral arteries is significantly lesser compared to that for the left arteries (p-value = 0.013 for left cerebral artery territories’s *cov*_*e*_: *cov*_*f*_ values being greater than for right). The left and right MCA are usually the bigger vessels amongst the six cerebral arteries, carrying a greater proportion of blood into the brain. Additionally, anatomically the MCAs have a greater tendency to receive emboli directly traversing the ICAs (e.g. as in Figure 4). As inertial embolic particles enter the aorta from the heart, they follow the greater curvature of the aortic arch, where they encounter the right branching vessels first, and follow on into the right carotid artery *(see supplementary animation)*. These factors together explain the observations on lower sensitivity of MCA territory cardiogenic embolus distribution, and stronger right brain preference, as observed from Figures 3 and 4. The prominent right-brain and MCA vasculature propensity of cardiogenic emboli have been reported in our prior studies [35] as well as clinical studies [24, 39], indicating that the reported trends here are consistent.

### 3.3 Circle of Willis anatomy potentially impacts proximal cervical flow

Our simulation framework incorporates the complete network of major supplying arteries from the heart to the CoW. This is advantageous for enabling investigations on correlations between flow at the cerebral arteries and at other locations upstream. Specifically, here we computed flow through the ICAs and BA (i.e. total flow through proximal cervical vessels), and compared them across the various patients and CoW anatomies considered. Figure 5, left panel, illustrates ratios between flow through L.ICA and BA (*Q_lica_*: *Q_ba_*; top), between R.ICA and BA (*Q_rica_*: *Q_ba_*; middle), and between R.ICA and L.ICA (*Q_rica_*: *Q_lica_*; bottom), for all the patient-anatomy combinations in our study. The corresponding panels on the right indicate how each of these ratios, for incomplete CoW anatomies, compare with corresponding ratios for CoW anatomy with a complete anastomoses. These relative ratio values on the right panels closer to unity indicate no influence of anatomical variants on proximal cervical flow. We observe that CoW anatomical variations induce significant changes in the proportion of flow across the ICAs and BA (p-values 0.003 and 0.006 for *Q_lica_*: *Q*_*ba*_ and *Q*_*rica*_: *Q*_*ba*_ values for incomplete CoW anatomies vs full CoW being different from 1.0, respectively), as well as induce noticeable but less significant variability in left-right asymmetry in ICA flow (p-value 0.7; *Q*_*lica*_: *Q*_*rica*_ between incomplete and complete CoW ≠ 1.0). Absence of LP1 and RP1 segments (fetal-type CoW anatomy) are observed to induce the greatest extent of variability in relative flow through the ICAs and BA, while absence of the AcoA, L.Comm, and R.Comm segments induce relatively smaller changes. We observe that absence of LP1 connector segment causes increased flow through left ICA with respect to BA, as well as reduces the right-left asymmetry in ICA flow (and vice-versa for absent RP1 connector segment). Additionally, in all cases, the fetal type variants lead to increased ipsilateral flow through the ICAs. The observed trends with respect to the fetal-type CoW anatomy variants can be explained as a consequence of ICA flow seeking to compensate for the deficit in PCA flow due to the BA not directly supplying the posterior cerebral vasculature. We note that these simulation results are based on the underlying assumption that the cerebral beds fed by the six major cerebral arteries maintain similar flow requirements across the various CoW anatomical variants. However, the findings are in agreement with previous clinical studies [45], indicating that CoW anatomy may influence how flow through the cervical vasculature reaches the brain.

**Figure 5:**
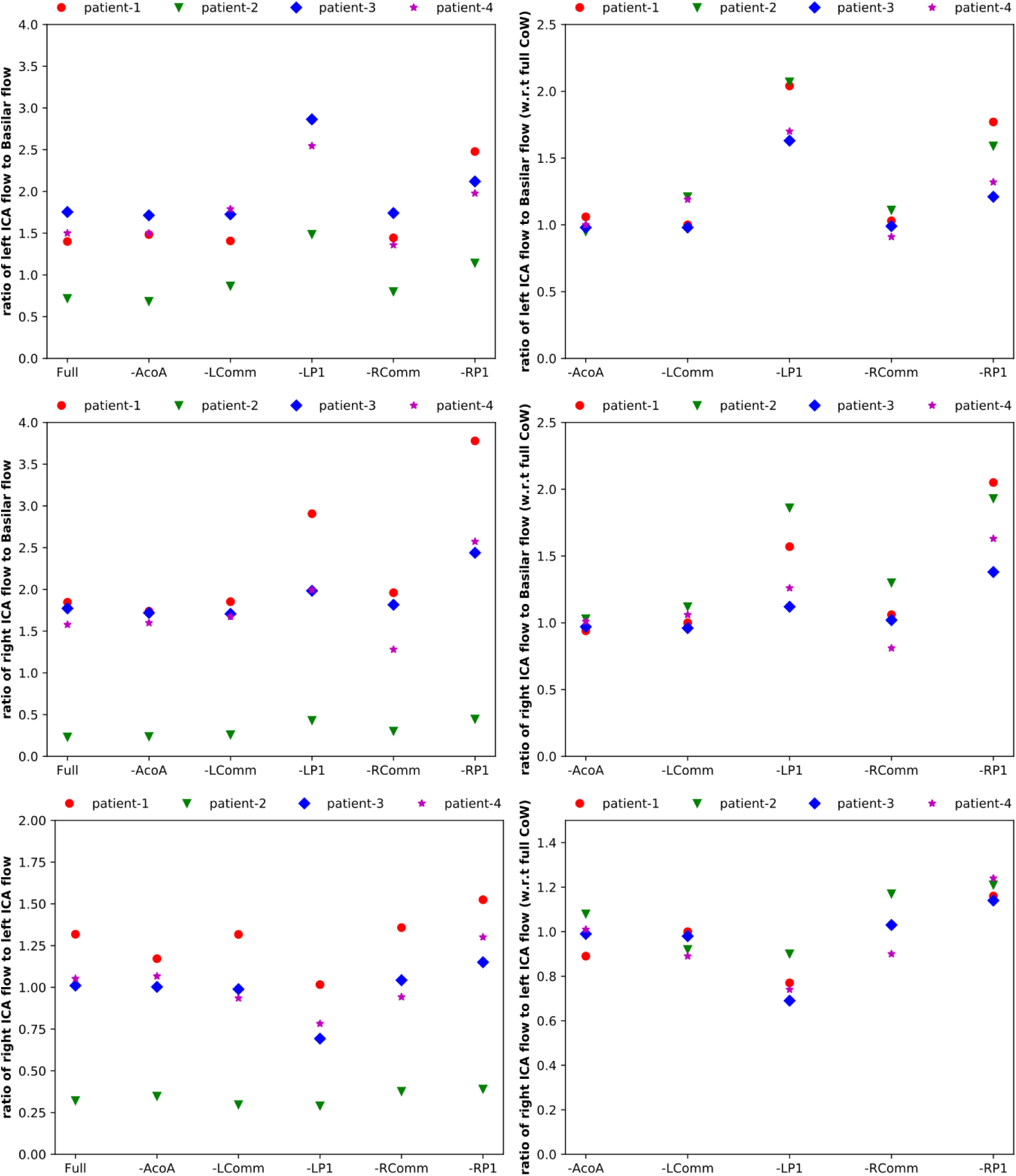
The ratios of incoming flow (into the brain) between (a) left ICA and BA (top,left); (b) right ICA and BA (mid,left); and (c) right and left ICAs (bottom,left); compared across all CoW anatomical variants considered, for all four patients. Corresponding panels on the right depict the respective flow-ratios relative to the models with a complete CoW. Supporting sample data for the flow-ratios included in supplementary figure S2.

### 3.4 Communicating arteries and their role in flow routing

Flow routing across the CoW communicating artery segments for all 24 patient-anatomy combinations was characterized by estimating volumetric flow across each communicating artery segment, using the method in Section 2.6. The results have been illustrated in Figures 6 and 7. Figure 6 depicts a map comprising the time-averaged volumetric flow-rate (over one cardiac cycle) across each communicating artery for the full CoW anatomy (the first column), and compares the factor by which the flow changes, as well as the flow direction, for all the remaining corresponding anatomical variations. We observe that CoW anatomy variations can alter not only the amount of flow routed across the communicating vessels, but also the direction in which flow is carried. The factors by which flow changes with respect to the full CoW variant (i.e. the ratios in columns 2-6 in Figure 6) are compiled in Figure 7. The left panel illustrates these flow-ratios classified based on CoW anatomy, while the right panel presents the ratios classified based on respective communicating vessels. Non-parametric Wilcoxon tests were conducted to establish which vessel/anatomy were associated with flow-ratios significantly different from unity. From the simulation results, the AcoA and L.Comm segments show the highest extent of variability in their flow engagement, as well as most significant extent of flow routing (respective p-values 0.004 and 0.002). Amongst the remaining vessels, R.Comm flow-ratios manifest high variability but not significant deviations from unity (p-value 0.2), while RP1 manifests low variability but significant deviations from unity (p-value 0.02). Contrarily, for CoW anatomies, the variants with missing LP1 and RP1 segments are seen to have significant differences in flow routed across the CoW (p-values 0.0014 and 0.0011 respectively) and also the highest extent of variability. Figure 6 indicates the L.Comm and R.Comm segments to be extensively engaged in flow distribution for CoW anatomy with missing LP1 and RP1. This is in contrast with all the other anastomoses where the proportion of flow carried by the L.Comm/R.Comm segments are quite low when compared to the left and right P1 and A1 connector segments. Our observations in Figure 6 and 7 also indicate that AcoA routes significant amount of flow and can reverse flow-direction, to maintain a left-right proximal collateral pathway for blood flow distribution in the brain. This quantitative evidence for the role of AcoA in the proximal collateral capacity of the CoW, is in agreement with observations from clinical studies on collateral flow characteristics of the brain [8, 25, 20, 49]. Furthermore, the general computational observations on CoW communicating arteries actively engaging in flow routing are in reasonable agreement with observations from clinical data [19].

**Figure 6:**
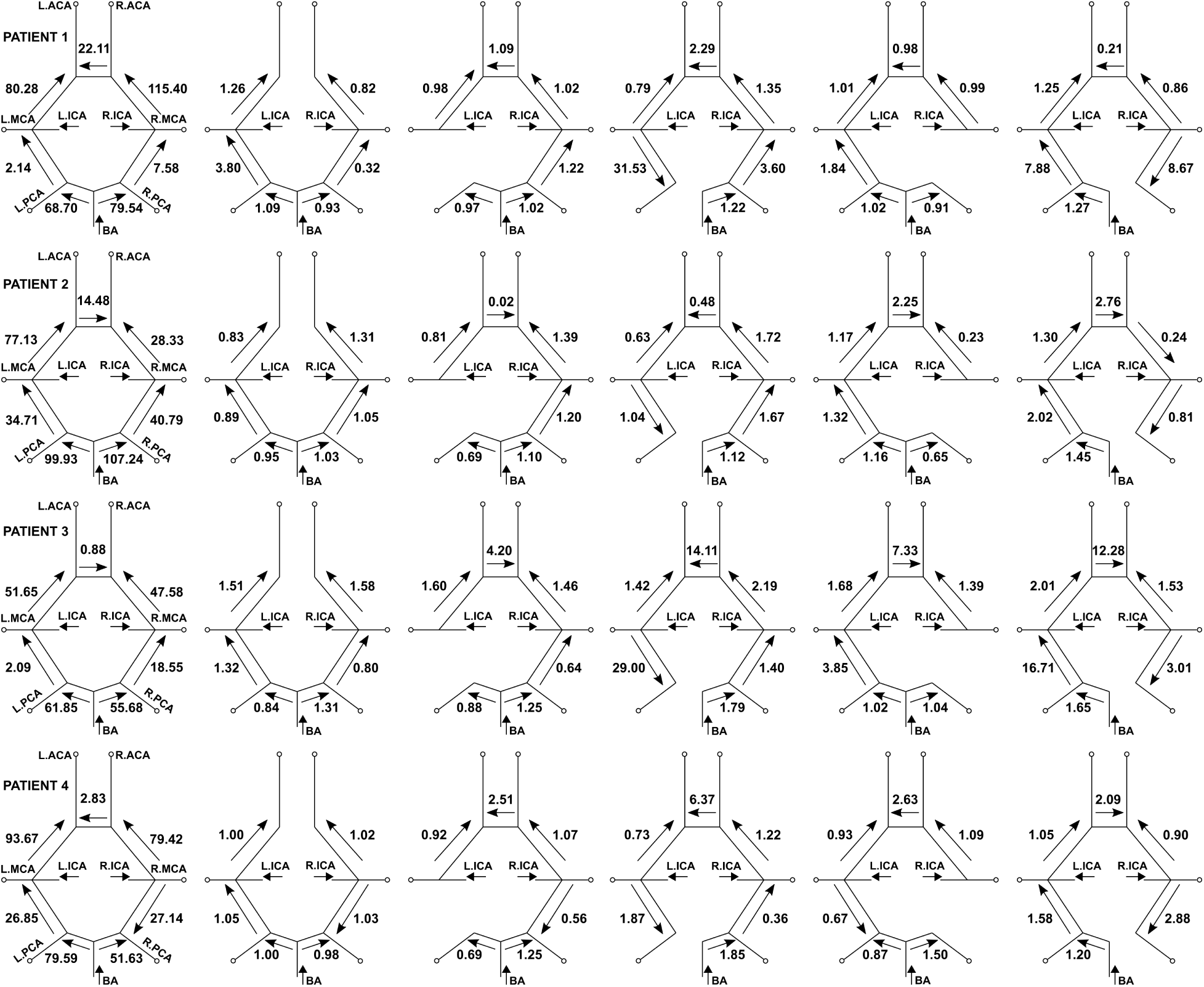
Map showing recruitment patterns for the various CoW communicating artery segments, compared across each anatomical variant. Arrows indicate direction of average flow. The first model in each row denotes the total average flow rate (ml/min) and flow direction. The remaining five anatomical variants are annotated based on the factor by which the average flow rate changes when compared with the corresponding flow for the full CoW model. Supporting flow-rate data included in supplementary figure S3.

**Figure 7:**
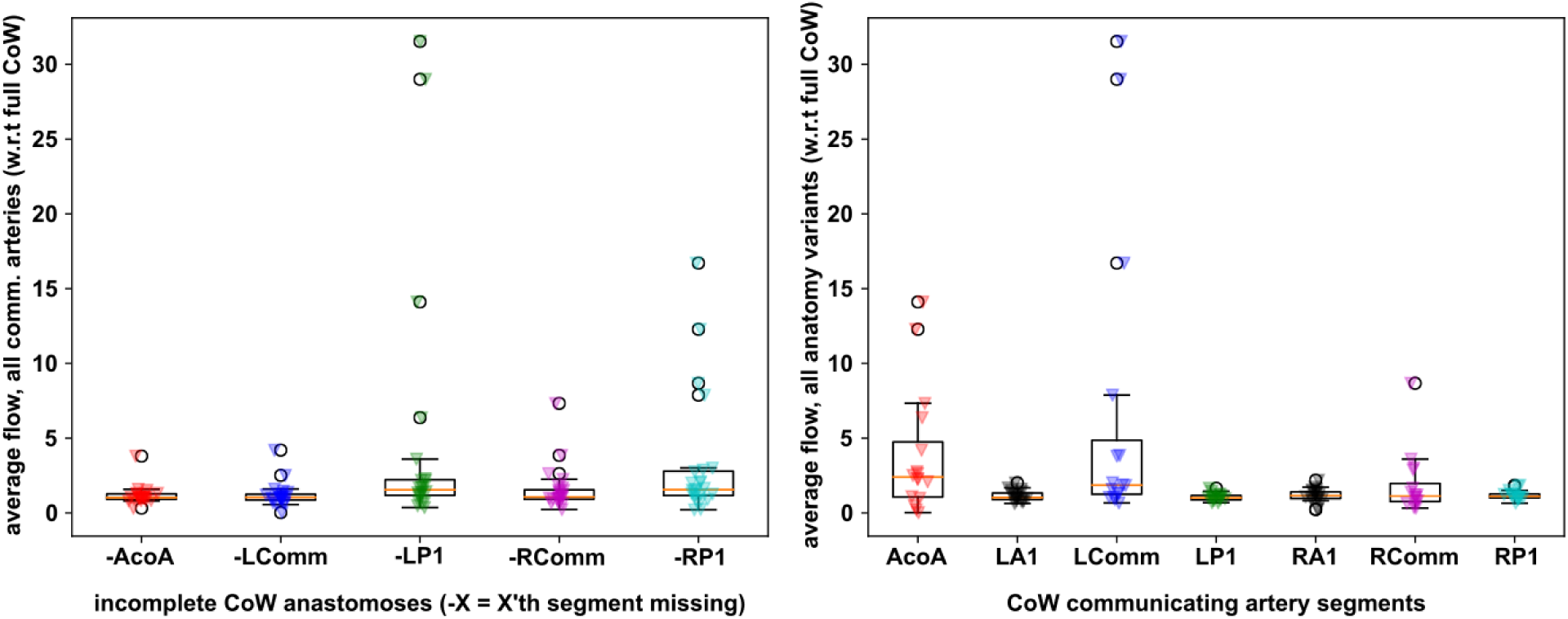
Quantification of flow routing for all the patient-CoW anatomy combinations. The left panel illustrates relative flow ratios with respect to full CoW cases, based on CoW anatomy. The right panel illustrates the same ratios based on communicating vessel segments. Value of unity, as in Figure 6, denotes no change in flow due to change in CoW anatomy. Supporting flow-rate data included in supplementary figure S3.

### 3.5 Embolus transport across communicating arteries

For elucidating the influence of CoW anatomy on embolus distribution, we further quantify embolus routing statistics using the routing count variable *R_k_* (discussed and defined in Section 2.6) for six communicating artery segments (indexed by *k*) - L.Comm, R.Comm, LA1, RA1, LP1, and RP1. Owing to the complexity and tortuosity of the anatomy at the location of the AcoA, it was not possible to get a clear definition of *R_k_* for this segment. The resultant *R_k_* for each communicating artery segment was compiled for all possible parameter combinations employed in this study, and post-processed to illustrate the trends in embolic particle transport across the CoW. First, *R_k_* values for each communicating segment in the incomplete CoW anatomies were compared with the corresponding values for the segments in the complete CoW, and the resulting ratios have been compiled in Figure 8.a. A value of 1.0 here would indicate that the number of emboli traversing a vessel did not change with CoW anatomy. The observed ratios in Figure 8.a. are significantly different from unity (p-value ≤ 0.05 for Wilcoxon test for ratios ≠ 1.0 for all vessels), indicating thereby that the resultant embolus trajectories, en-route to exiting one of the six major cerebral arteries, are significantly influenced by the CoW anatomy variations. The relative importance of each communicating artery segment in transporting emboli is illustrated in Figure 8.b., where *R*_*k*_ values for each of the arteries within a particular CoW anatomy have been normalized by the maximum *R*_*k*_ observed for that anatomy. Values closer to unity in Figure 8.b. indicate a high extent of embolus traversal across the vessel segment and vice-versa. The results indicate that the LA1 and RA1 segments are associated with the greatest extent of embolus traversal, and the L.Comm and R.Comm segments are associated with the least extent of embolus traversal. Influence of missing vessels is illustrated in Figure 8.c., where maximum *R*_*k*_ values for thrombo-emboli, relative to the same values for the full CoW model, is categorized based on CoW anatomy. Samples greater than unity denote higher extent of embolus routing, and the most significant extent of routing is observed for anatomies with missing LP1 and RP1 segments (p value 0.0016 and 0.017 respectively for relative *R*_*k*_ ≠ 1.0). This follows the trend observed in Section 3.4 for flow routing. The influence of the individual embolus properties (size, and material composition) have been isolated and illustrated in Figure 8.d, which depicts the total *R*_*k*_ values across all the different communicating vessels respectively. A high value along the y-axis denotes that a large number of emboli were routed across the vessels, while a low value indicates minimal engagement of the vessels in distributing emboli to the distal cerebral vasculature beds. The results indicate a strong influence of the embolus particle inertia (illustrated also in Figure 4), as well as density ratio between embolus and blood, in governing their re-routing across the CoW - with the general trend being that larger and denser emboli tend to enter and exit the CoW with less traversal across the communicating vessels.

**Figure 8:**
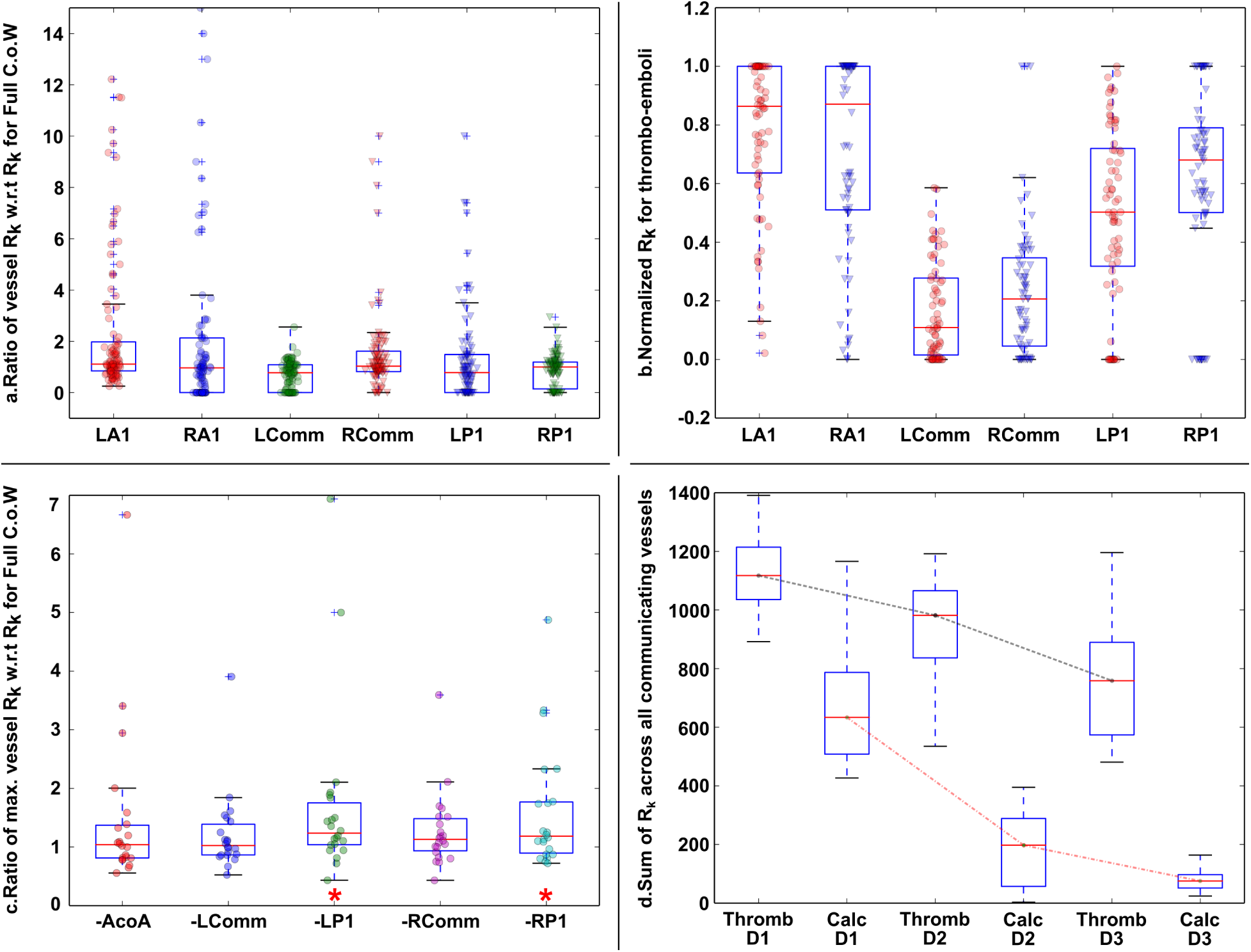
Illustration of the results for embolus routing analysis. Panel a. depicts the ratio between routing count (R_k_) through each vessel in a model, compared with that of the same vessel for the full CoW model. Panel b. depicts the sample statistics for R_k_ for each vessel within a CoW anatomy normalized with respect to the maximum R_k_ observed for that anatomy. Corresponding data for all emboli included in supplementary figure S4. Panel c. depicts variation in extent of embolus traversal by categorizing maximum R_k_ across each CoW anatomy (* denotes segments with ratios differing from unity with p-value ≤ 0.05). Panel d. depict the variation in the extent of embolus traversal across the CoW, quantified by the total R_k_ values for each embolus type, with respect to embolus size and material composition. The two dashed lines indicate different material composition, and variations along each reflects the role of momentum response time of emboli. Additional data on embolus routing is included in supplementary figures S5 & S6.

## 4 Discussion

### 4.1 Cerebral vasculature and cardio-embolic stroke

Our simulations illustrate the routing of flow across the CoW artery segments for varying anatomy, and indicate that interaction of inertial embolic particles with re-routed flow leads to differences in embolus distribution to the major cerebral vascular beds. Consequently, cerebral vasculature can be thought to play a bipartite role in ischemic events. The first, and better known, role is in terms of flow compensation during occlusion, and collateral circulation. The second role, as demonstrated here, comprises the cerebral vasculature influencing embolus transport leading to potential ischemia. Mechanistically, emboli entering the CoW from a supplying vessel (e.g. ICA) may not exit into the immediately connected cerebral artery (e.g. MCA) as expected from anatomical considerations alone, but instead traverse several vessel segments to exit at another location *(see supplementary animation)*. These insights on the role of cerebral vasculature on embolic stroke obtained from our study are critical for evaluating the source of emboli [44]. Thus, our model would allow clinicians to direct therapy (surgical, interventional or pharmacologic) to the appropriate anatomy in patients with multiple potential sources of emboli. Resolving embolus source is particularly important in patients with ambiguous clinical history such as moderate diffuse atherosclerosis with a hypoperfusive event. Also of clinical relevance are cases of artery to artery microemboli in intracranial atherosclerosis or other arterial disease [47, 13]. Our data suggest that these microemboli trajectories are more influenced by CoW anatomy and thus may generate non-intuitive locations for distal infarction.

### 4.2 The triad of anatomy, flow, and embolus properties

The biomechanical events governing embolus transport to the brain comprise a synergistic interplay of anatomical features, hemodynamics, and embolus properties. Anatomical features like vessel curvature and tortuosity, vessel segment interconnectivity, and branching has a strong influence on the three-dimensional flow patterns through these vessels. Pulsatile viscous flow through realistic anatomical vasculature is associated with extensive vorticity, helical flow, swirling flow, and retrograde flow during the deceleration phase of the cardiac cycle. Embolus particles of finite size and inertia interact with these spatio-temporally varying flow features, leading to a nearly chaotic advection from their origin site to their destination [37] *(see supplementary animation)*. This anatomy-flow-embolus synergy can be utilized to devise tools that integrate quantitative vascular geometry analysis [12] with embolus dynamics [36]. This can enable embolism predictions based on medical image-based vascular geometry information, which may have diagnostic utility owing to geometry analysis times being comparable with clinical timings. Furthermore, mathematical analysis of the underlying equations governing embolus dynamics reveals two key non-dimensional variables that govern their motion [36]. First, the momentum response time (often referred by the related non-dimensional Stokes number) governs how quickly the particle responds to flow-induced forces. Second, the density ratio between the embolus and blood, which governs how buoyant the embolus is within the vessel. This is clearly observed in Figure 8.d., where the two dashed lines represent constant density-ratios, and variations along each represent how momentum response time influences *R*_*k*_. In prior work, we have isolated the influence of these three factors in the context of anatomy of the aortic arch, determining whether emboli are routed towards or away from the brain. The results from this study illustrates the same influence in the context of CoW anatomy that determines which of the six cerebral vasculature beds does an incoming embolus go to.

### 4.3 Assumptions and limitations

The underlying computational modeling framework was based on few key assumptions. First, as discussed in our prior works, the influence of the embolus on blood flow was not accounted for, leading to a one-way interaction between embolus and flow. Hence, our study focused on emboli sizes in the small-to-medium regime with respect to the vessel diameters, such that potential occlusions in vasculature beds at or proximal to the M1, A1, and P1 segments were not considered. While a fully-resolved coupling is mathematically more accurate, it is also significantly expensive, which would render the kind of sampling-based analysis presented here untenable. The one-way interaction model used here has been validated previously for industrial and cardiovascular applications. Our prior work demonstrated that this approach is reasonable for the small-to-medium size emboli considered here [36]. Further in-depth investigation on fully resolved two-way fluid particle interactions for complex cardiovascular flows is an area that needs separate investigation by itself, and comprises a parallel focus of our current research interests. Second, we assumed that removal of one communicating vessel segment from the CoW would not influence the size and length of the remaining segments. This was primarily because well characterized human population data on this was not available, and introducing arbitrary geometric rules to model any vessel size changes was avoided in our framework. However, our image-based modeling procedure is flexible in dealing with individual vessels at the level of image-segmentations, and any trends in vessel shape with respect to CoW topology can be easily incorporated. Third, the influence of cerebral autoregulation has not been included within the model. While this may influence model predictions for embolus transport in patients who have already had a stroke or transient ischemic attack, this was not a factor for our study as we considered healthy patients, and focused on the event horizon prior to any cerebro-embolic event. On a related note, boundary conditions at the inlet and cerebral outlets were kept consistent across patient models here, in accordance with the controlled multi-factor experiment design (Section 2.5). However, the framework poses no inherent complications in employing patient-specific boundary conditions from imaging [34] or other modalities, which can enable targeted embolus dynamics predictions for subjects. Finally, a key study limitation was the small population size considered here. While we considered four different patients with six different CoW topological variants, this was still a small subject cohort size, and renders some restrictions in terms of characterizing inter-patient variabilities, patient statistics, and influence of hypoplasia/aplasia in particular CoW artery segments. The findings presented here, however, retain their significance in terms of providing numerical evidence on the role of CoW anatomy in embolus transport and embolic stroke risk, and the corresponding hemodynamic connection.

### 4.4 Broader implications

The computer simulation framework and data, described here, have broader implications in terms of stroke diagnosis and treatment. Our approach can complement current clinical imaging protocols in enabling patient-specific predictions on embolic risks by integrating CT/MR/Trans-cranial Doppler data with numerical modeling of various embolism scenarios. This can be used to disambiguate embolus source in patients with multiple possible sources. Additionally, our results show that likelihood of cardio-embolic sources being responsible for specific infarct patterns comprises both general patterns and patient specific patterns. One notable general pattern is the right MCA territory being least variable in its likelihood of receiving emboli. It remains intriguing that our data suggest a protective effect for the left hemisphere vis a vis embolic material; indicating that anatomy and physiology apparently recognize the importance of language dominance and the devastating effects of left MCA stroke. Furthermore, insertion of catheters or other endovascular devices commonly employed in surgical treatments, can effectively alter blood flow in similar ways as hypoplasia/aplasia of communicating arteries in the CoW [30]. Thus, collateral capacity through communicating arteries of the CoW influence ischemic risks during procedures like carotid endarterectomy [23]. The methodology developed here can help address effectiveness of cerebral vasculature in maintaining perfusion during a procedure, and evaluating peri-operative embolic stroke risks due to emboli released during a procedure due to endothelial trauma [1], or plaque rupture. Finally, the influence of CoW anatomy on flow re-distribution and particle routing, is also important for drug delivery techniques in the brain. For example, shear activated particle-based drugs for thrombolysis have shown promise in recent investigations [26, 32]. The tools and findings described here could be beneficial in successfully transforming such techniques into more effective stroke treatment approaches.

### 4.5 Concluding Remarks

We have described a patient-specific simulation based study to investigate the role of the Circle of Willis (CoW) anatomy in cardio-embolic stroke. We performed systematic parametric experimentation on transport of cardiogenic emboli of varying sizes and compositions in four patient vasculature geometries, considering six different CoW anatomical variants. The resultant data and statistics helped establish that CoW anatomical variations play a role in altering embolus distribution in the brain, and that MCA territory embolism risks are least sensitive to this anatomical influence. The ability to extract fully resolved flow and embolus trajectory information also helped illustrate the mechanical phenomena that possibly leads to these variations in embolus distribution. Our findings indicated that CoW anatomical variations lead to differences in flow routing from the proximal cervical vessels, across the various communicating artery segments of the CoW, into the six major cerebral arteries. Embolic particles of finite size and inertia interact with the flow routed across the vessel segments, and traverse the CoW based on these interactions, leading to a difference in their final destination cerebral vasculature. The computational approach as well as the resulting findings have broad impacts on understanding stroke biomechanics, stroke diagnosis, and treatment planning.

## 6 Acknowledgements

This work was supported by the American Heart Association Award: 13GRNT17070095. This research used the Savio computational cluster resource provided by the Berkeley Research Computing program at the University of California, Berkeley. NDJ acknowledges support from the Regent’s and Chancellor’s Research Fellowship at U.C. Berkeley. DM, NDJ, and SCS conceptualized the design of the study. DM developed the computational framework, performed the embolus dynamics, performed all statistical and data analysis, drafted the manuscript. NDJ devised the image-based modeling framework, computed all flow simulations, and contributed to embolus dynamics simulations. JN helped with data analysis, and contributed clinical and diagnostic connections to the simulation data. NDJ, SCS, and JN reviewed and edited the manuscript draft. Final manuscript version was in agreement with all Authors.

## Supplementary Material

### Animation of embolus dynamics

A supplementary video file has been included to help visualize various features of the embolus dynamics across the arterial network. The animation illustrates the influence of embolus size and embolus material composition on their dynamics. Additionally, a zoomed view of the embolus dynamics within the Circle of Willis for one of the patient-anatomy models has been included to illustrate their movement across the various segments of the Circle.

### Supplementary data and figures

We have included here a collection of figures and processed quantitative data which support and supplement the text and the figures presented in the main manuscript. For each supplementary figure/table included here, the relevant descriptions are presented in the captions. Respective figures have been referenced at appropriate locations within the main text.

**Table S1:** Simulation data for coefficients of variation of thrombo-emboli distribution to the six major cerebral arteries across all anatomical variations considered, and the ratio between embolus and flow coefficients of variation. Varying embolus sizes denoted by 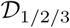.

| Cases |  | Cerebral arteries |  |  |  |  |  |
| --- | --- | --- | --- | --- | --- | --- | --- |
| Patient | Size | LACA | LMCA | LPCA | RACA | RMCA | RPCA |
| Patient: $\mathcal{P}_1$ | Ratio of $(c.o.v)_e$ w.r.t. $(c.o.v)_f$ | | | | | | |
| | $\mathcal{D}_1$ | 2.59 | 0.76 | 2.95 | 3.42 | 0.85 | 1.55 |
| | $\mathcal{D}_2$ | 1.04 | 0.58 | 5.11 | 6.02 | 0.55 | 2.43 |
| | $\mathcal{D}_3$ | 4.02 | 1.03 | 6.08 | 7.98 | 1.43 | 2.93 |
|  | Sample coefficients of variation |  |  |  |  |  |  |
| | $\mathcal{D}_1$ | 0.0798 | 0.0761 | 0.0723 | 0.0713 | 0.0810 | 0.0964 |
| | $\mathcal{D}_2$ | 0.0320 | 0.0584 | 0.1251 | 0.1254 | 0.0530 | 0.1513 |
| | $\mathcal{D}_3$ | 0.1241 | 0.1032 | 0.1488 | 0.1662 | 0.1366 | 0.1824 |
|  | flow | 0.0309 | 0.1006 | 0.0245 | 0.0208 | 0.0956 | 0.0622 |
| Patient: $\mathcal{P}_2$ | Ratio of $(c.o.v)_e$ w.r.t. $(c.o.v)_f$ | | | | | | |
| | $\mathcal{D}_1$ | 6.07 | 1.16 | 1.03 | 1.95 | 0.90 | 1.20 |
| | $\mathcal{D}_2$ | 7.46 | 1.10 | 1.97 | 2.63 | 0.51 | 1.41 |
| | $\mathcal{D}_3$ | 43.16 | 2.39 | 4.30 | 8.70 | 0.91 | 3.33 |
|  | Sample coefficients of variation |  |  |  |  |  |  |
| | $\mathcal{D}_1$ | 0.1333 | 0.1744 | 0.2013 | 0.2106 | 0.2036 | 0.2613 |
| | $\mathcal{D}_2$ | 0.1637 | 0.1663 | 0.3833 | 0.2845 | 0.1159 | 0.3065 |
| | $\mathcal{D}_3$ | 0.9470 | 0.3597 | 0.8374 | 0.9394 | 0.2056 | 0.7256 |
|  | flow | 0.0219 | 0.1506 | 0.1947 | 0.1080 | 0.2255 | 0.2180 |
| Patient: $\mathcal{P}_3$ | Ratio of $(c.o.v)_e$ w.r.t. $(c.o.v)_f$ | | | | | | |
| | $\mathcal{D}_1$ | 1.10 | 1.05 | 1.84 | 0.96 | 0.90 | 0.99 |
| | $\mathcal{D}_2$ | 1.50 | 1.70 | 2.67 | 0.76 | 0.79 | 1.23 |
| | $\mathcal{D}_3$ | 3.65 | 4.03 | 7.34 | 1.83 | 1.08 | 1.30 |
|  | Sample coefficients of variation |  |  |  |  |  |  |
| | $\mathcal{D}_1$ | 0.1786 | 0.0876 | 0.1437 | 0.1614 | 0.1673 | 0.1785 |
| | $\mathcal{D}_2$ | 0.2437 | 0.1416 | 0.2088 | 0.1282 | 0.1479 | 0.2218 |
| | $\mathcal{D}_3$ | 0.5922 | 0.3364 | 0.5734 | 0.3080 | 0.2009 | 0.2348 |
|  | flow | 0.1621 | 0.0835 | 0.0782 | 0.1680 | 0.1869 | 0.1808 |
| Patient: $\mathcal{P}_4$ | Ratio of $(c.o.v)_e$ w.r.t. $(c.o.v)_f$ | | | | | | |
| | $\mathcal{D}_1$ | 5.97 | 2.21 | 2.94 | 5.47 | 4.27 | 2.34 |
| | $\mathcal{D}_2$ | 5.91 | 2.52 | 4.28 | 12.80 | 3.75 | 2.23 |
| | $\mathcal{D}_3$ | 47.31 | 2.13 | 18.75 | 18.93 | 4.36 | 3.50 |
|  | Sample coefficients of variation |  |  |  |  |  |  |
| | $\mathcal{D}_1$ | 0.0842 | 0.0421 | 0.0945 | 0.0545 | 0.0913 | 0.1059 |
| | $\mathcal{D}_2$ | 0.0833 | 0.0481 | 0.1377 | 0.1275 | 0.0801 | 0.1011 |
| | $\mathcal{D}_3$ | 0.6668 | 0.0406 | 0.6036 | 0.1885 | 0.0932 | 0.1584 |
|  | flow | 0.0141 | 0.0191 | 0.0322 | 0.0100 | 0.0214 | 0.0452 |

**Table S2:**
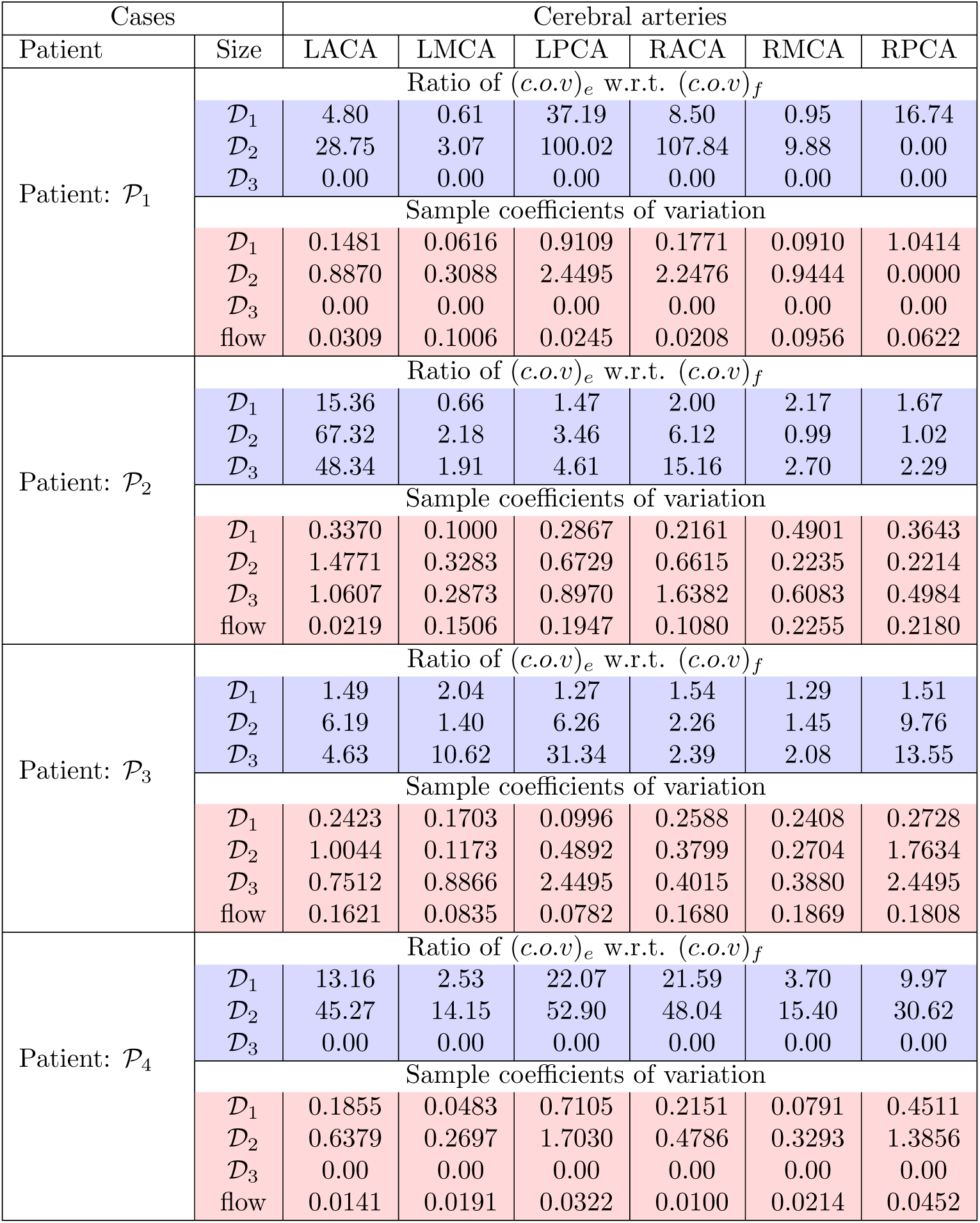
Simulation data for coefficients of variation of calcified emboli distribution to the six major cerebral arteries across all anatomical variations considered, and the ratio between embolus and flow coefficients of variation. Varying embolus sizes denoted by 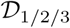. Note that for patients 1 & 4 largest calcified embolic fragments are too heavy to reach the brain at all.

**Figure S1:**
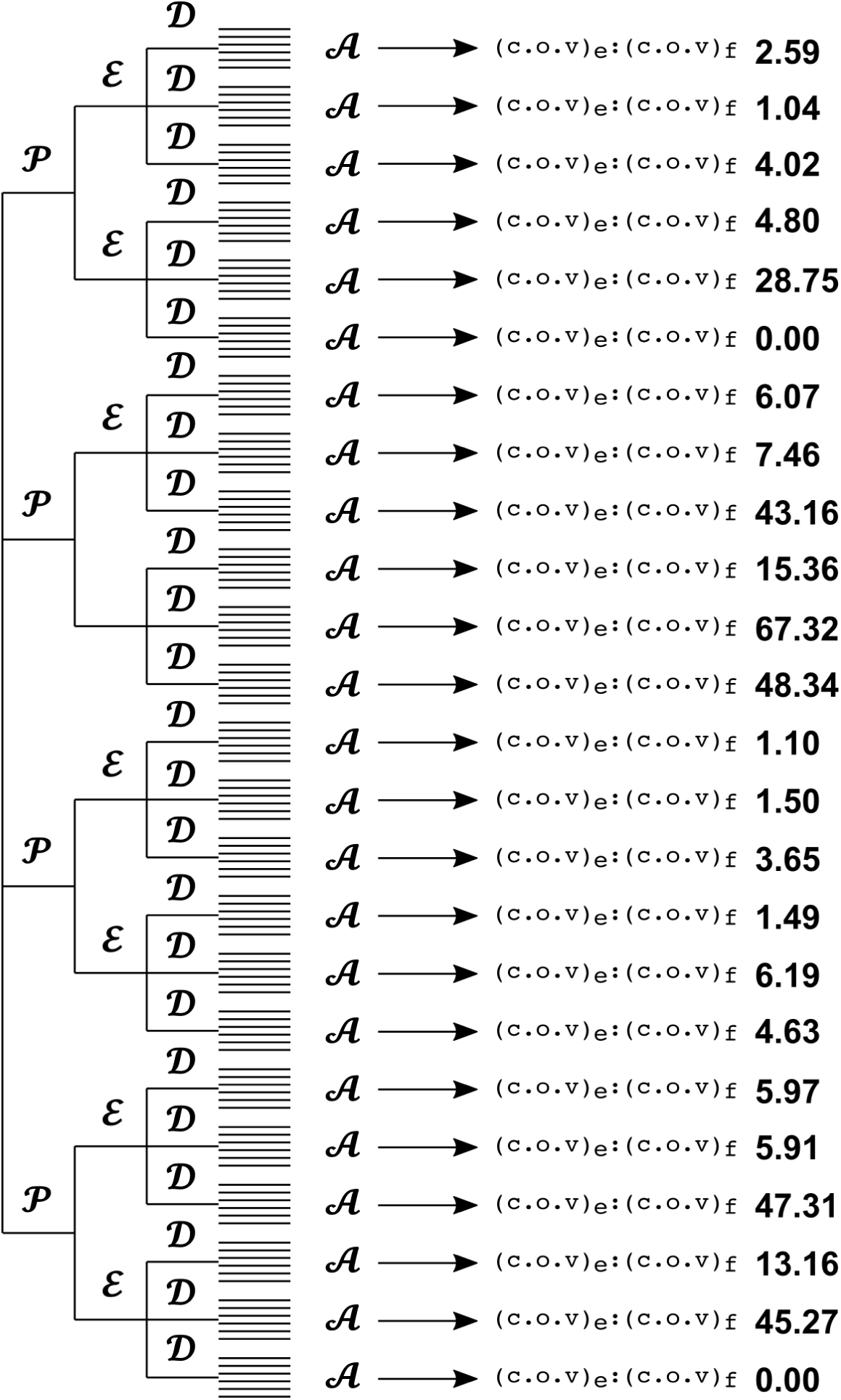
Supporting figure providing a visual depiction of the parametric experimentation set-up and the corresponding variability quantification (using cov_e_: cov_f_). Numerical data for L. ACA embolus distribution have been shown here. Figure supplements the explanation of the data in Tables S1 and S2 above, and Table 2, and Figure 3 in the article.

**Figure S2:**
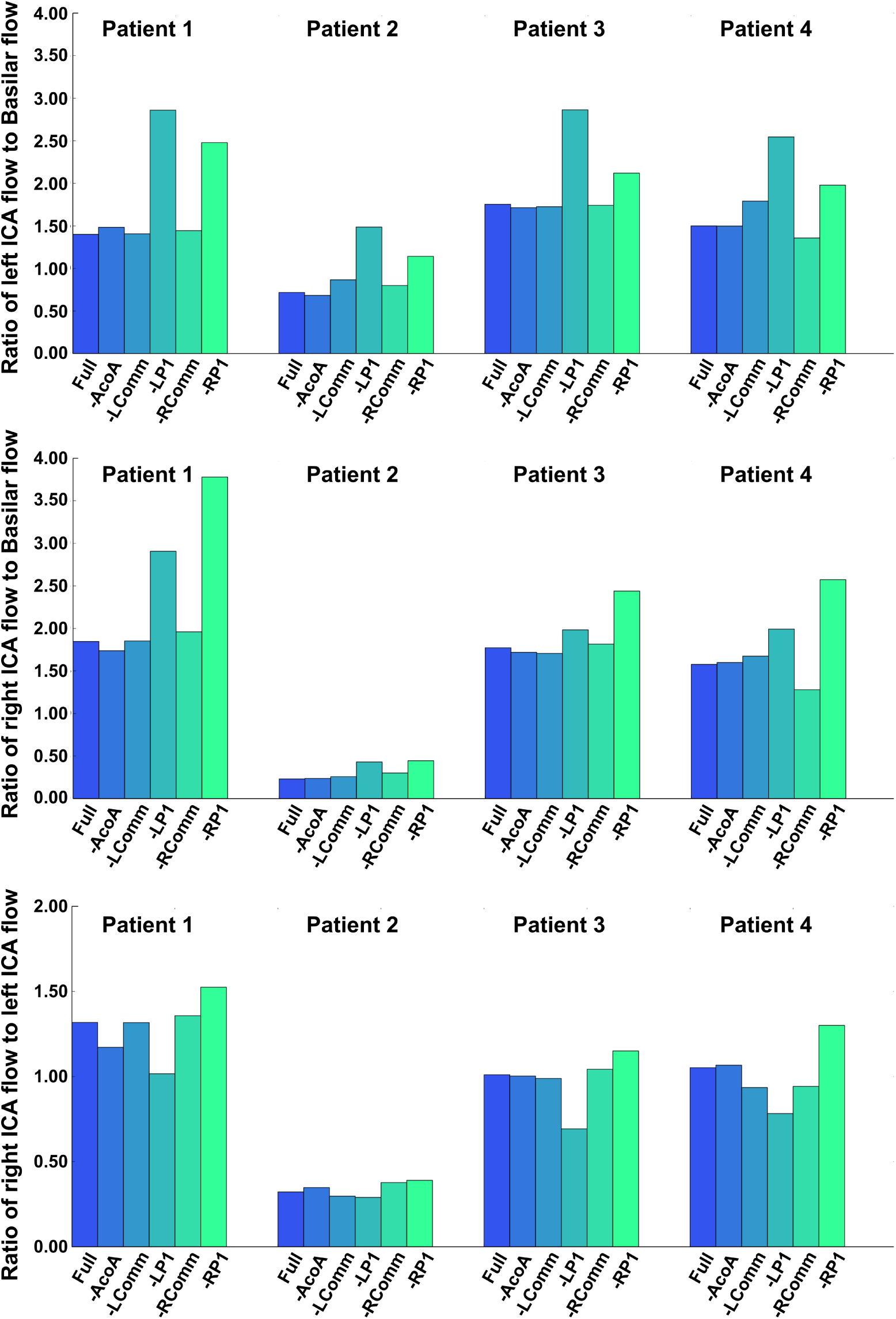
Visualizing the data for the relative incoming flow (into the brain) between (a) left internal carotid and basilar arteries (top); (b) right internal carotid and basilar arteries (mid); and (c) right and left internal carotid arteries (bottom); compared with respect to patient and Circle of Willis anatomical variants considered. Figure supplements the visualization presented in Figure 5 in the article.

**Figure S3:**
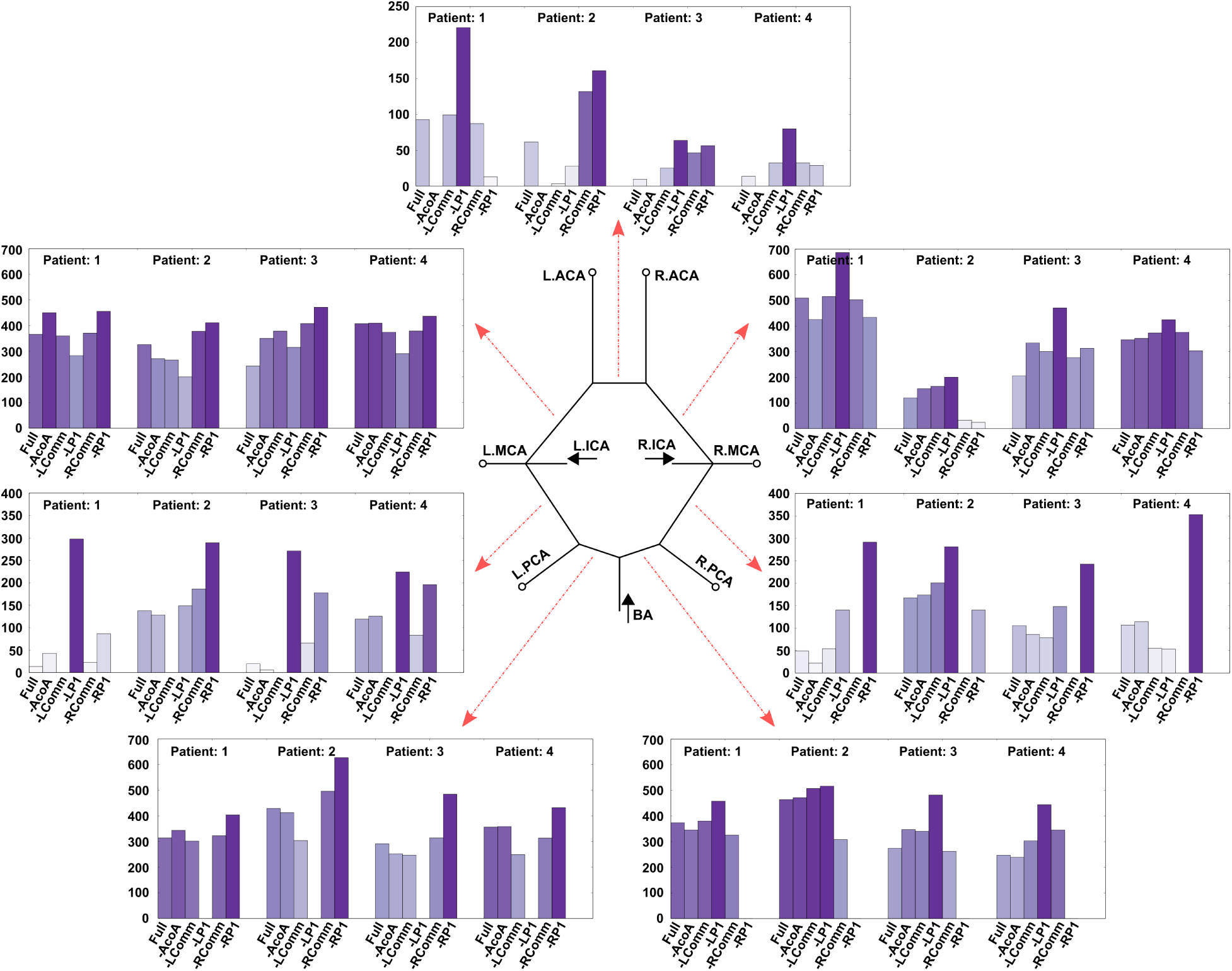
Visualization of data for peak systolic flow through each communicating artery segment, compared across all Circle of willis anatomical variations, for each patient. The dotted lines indicate the artery segments that were absent in the anastomoses considered. Figure supplements the visualization presented in Figure 7 in the article. All flow-rates are in ml/min.

**Figure S4:**
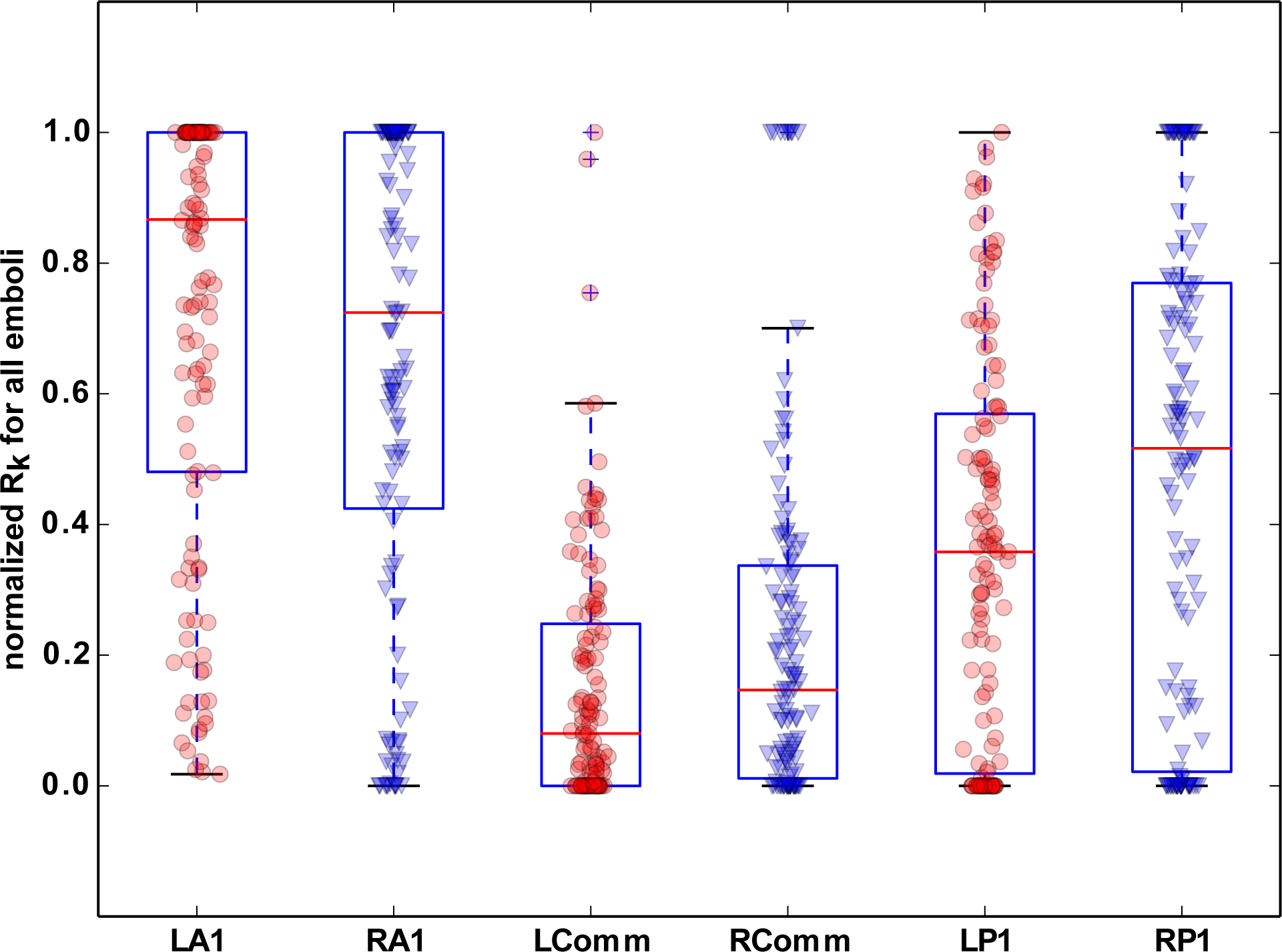
Normalized routing count (R_k_) values compiled for all the embolus sizes and material compositions considered, computed across all the Circle of Willis anatomical variations.

**Figure S5:**
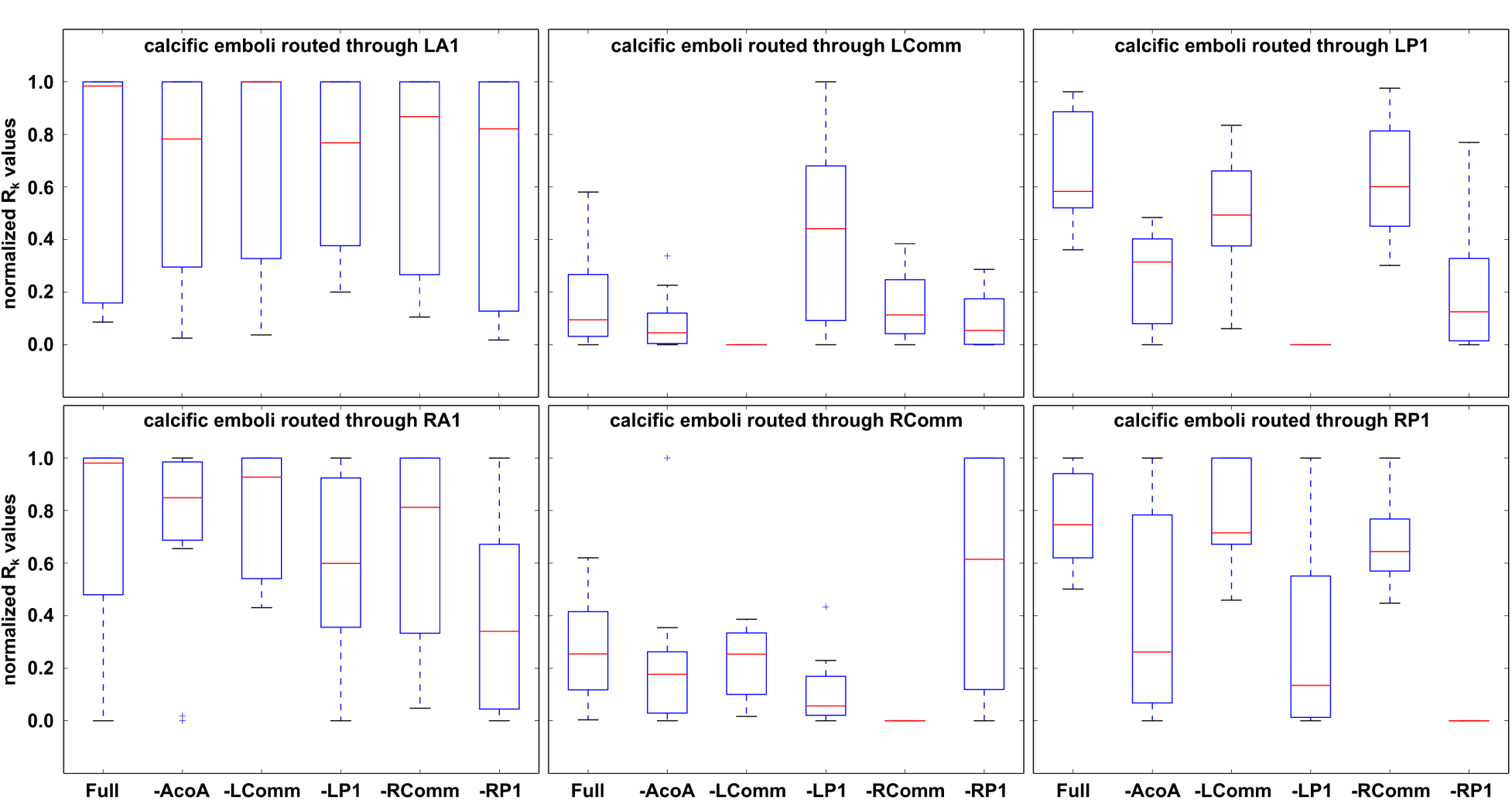
Description of thrombo-embolus routing through the Circle of Willis for varying anastomoses, illustrated for each communicating artery segment (except the anterior communicating artery as described in Section 3.5).

**Figure S6:**
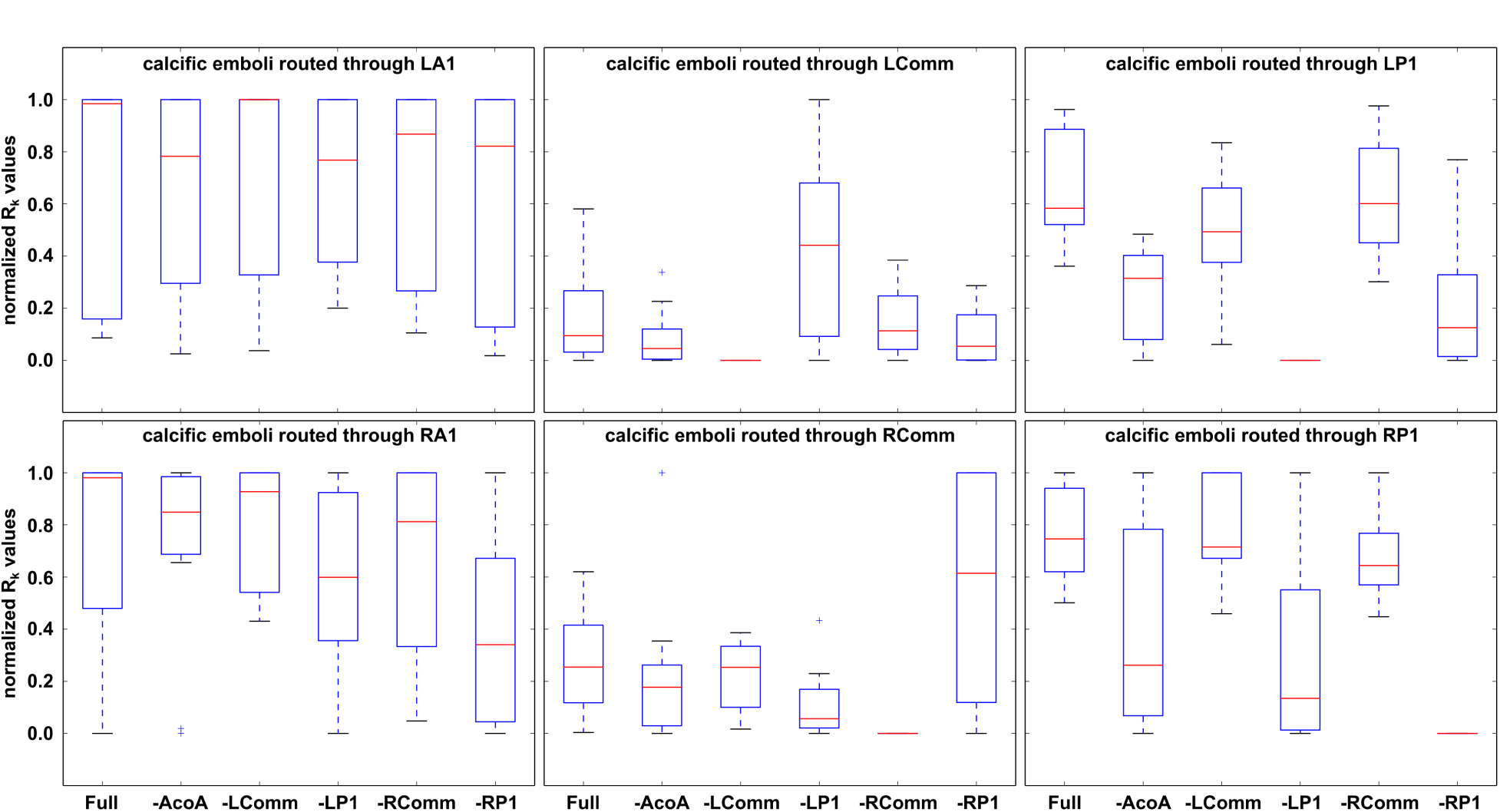
Description of cacific-embolus routing through the Circle of Willis for varying anastomoses, illustrated for each communicating artery segment (except the anterior communicating artery as described in Section 3.5)

